# Shifts in Naturalistic Behaviors Induced by Early Social Isolation Stress are Associated with Adult Binge-like Eating in Female Rats

**DOI:** 10.1101/2024.10.28.620673

**Authors:** Timothy B. Simon, Julio Sierra, Arianna Williams, Giara Wright, Allison Rhee, Julius Horn, John Lou, Fransua Sharafeddin, Perla Ontiveros-Angel, Johnny D. Figueroa

## Abstract

Binge eating (BE) is a highly pervasive maladaptive coping strategy in response to severe early life stress such as emotional and social neglect. BE is described as repeated episodes of uncontrolled eating and is tightly linked with comorbid mental health concerns. Despite social stressors occurring at a young age, the onset of BE typically does not occur until adulthood providing an interval for potential therapeutic intervention. Currently, our knowledge of longitudinal noninvasive digital biomarkers predictive of BE needs further development. Monitoring longitudinal impacts of adolescent social isolation stress on naturalistic behaviors in rats will enable the identification of noninvasive digital markers of disease progression to predict adult eating strategies. Recognizing adolescent naturalistic behaviors shaped by social stress informs our understanding of the underlying neurocircuits most effected. This study aimed to monitor and identify longitudinal behavioral shifts to enhance predictive capabilities in a rat model of social isolation stress-induced BE. We placed Paired (n=12) and Socially Isolated (SI, n=12) female rats in observational home cages weekly for seven weeks to evaluate the effect of SI on 10 naturalistic behaviors. All 10 naturalistic behaviors were simultaneously detected and tracked using Noldus Ethovision XT automated recognition software. Composite phenotypic z-scores were calculated by standardizing all 10 behaviors. When transitioning into adulthood, all rats underwent conventional emotionality testing and were exposed to a Western-like high fat diet (WD, 43% kcal from fat) to evaluate BE. Longitudinal assessments revealed SI-induced shifts in adolescent phenotypic z-scores and that sniffing, unsupported rearing, jumping, and twitching were the most susceptible to SI. SI increased emotionality compared to the Paired controls. Finally, we identified adolescent twitching as a digital biomarker of adult WD consumption. This study employed novel approaches to identify early life predictive biomarkers of adult eating strategies and laid a foundation for future investigations.

## INTRODUCTION

Eating disorders are serious, chronic conditions that represent major public health issues with an estimated 17.3 million globally recorded cases (Santomauro et al., 2021). In the United States, binge eating disorder is the most common eating disorder with adult women (1.25%) being three times more likely to develop the disorder than men (0.42%) (Hudson et al., 2007; Udo and Grilo, 2018). Despite the underlying causes of binge eating being multifactorial, psychosocial distress during childhood or adolescence is a major contributing factor (Brewerton, 2022; Chu et al., 2022; Cortés García et al., 2022; Klatzkin et al., 2024; Quilliot et al., 2019; Shin and Miller, 2012; Talbot et al., 2013; Wattick et al., 2023). Adolescents can develop an eating disorder, but the average age of disease onset is not until 25 years old making it difficult to pinpoint how adolescent psychosocial stress contributes to disease ontology (Hudson et al., 2007; Udo and Grilo, 2018). Most preclinical research has traditionally focused on stable "trait-like" differences—such as impulsivity, locomotion, and anxiety-like behaviors—as key characteristics of rats prone to maladaptively cope with excessive or unhealthy eating (Boggiano, 2013; Corwin, 2004; Karatayev et al., 2010; Rodríguez-Rangel et al., 2024; Velázquez-Sánchez et al., 2014). However, observing unperturbed naturalistic behaviors in the same individual through time may better inform our understanding of underlying neurocircuits and molecular mechanisms generating adverse coping strategies and ultimately aid in enhancing predictive measures (Ashby et al., 2010; Boggiano et al., 2005; Rodríguez-Rangel et al., 2024).

Social stress, particularly emotional neglect during the neurodevelopmentally vulnerable stage of adolescence, is closely linked to the development and exacerbation of eating disorders (Batista, 2018; Cortés García et al., 2022; Klatzkin et al., 2024). More than 60% of adults have experienced early life social stress developing long term consequences persisting into adulthood (Agorastos et al., 2019; Swedo et al., 2023). Stressful social situations, such as perceived loneliness and social isolation, can trigger maladaptive coping mechanisms, including disordered eating behaviors (Chu et al., 2022; Herhaus et al., 2020; Makri et al., 2022; Quilliot et al., 2019; Sailer et al., 2022; Tremolada et al., 2016). Individuals experiencing high levels of social stress may seek comfort food, leading to binge eating in an attempt to exert control over their environment (Dallman et al., 2005; Klatzkin et al., 2024; Montigny et al., 2013). Adverse environmental insults alter brain function and hormone levels, exacerbating disordered eating (Fowler et al., 2019; Gluck et al., 2004; Goldfield et al., 2008; Vega-Torres et al., 2022, 2018). Investigations in rodents, particularly rats, have revealed the impact of social isolation stress on behavior and neuronal health during adolescence (Gądek-Michalska et al., 2017; Weiss et al., 2004; Yorgason et al., 2016). Rats raised in isolation exhibit behavioral abnormalities, such as elevated anxiety-like and fear conditioned behavior and dampened acoustic startle responsivity (Weiss et al., 2004); altered and elevated peripheral cytokine release (Gądek-Michalska et al., 2017); modified hippocampal synaptic plasticity (Djordjevic et al., 2012); enhanced glutamatergic neurotransmission while decreasing AMPA receptors in CA1 pyramidal neurons (Gómez-Galán et al., 2016); and altered central dopaminergic systems (Yorgason et al., 2016), thus mirroring phenotypes observed in human psychiatric disorders. The pronounced connection between social stress and eating disorders highlights the importance of addressing social factors in prevention and treatment strategies. These findings suggest that early life social environments play a crucial role in shaping eating behaviors and contribute to the emergence of disordered eating patterns in adolescence and adulthood. Understanding the molecular underpinnings of this connection may provide new therapeutic avenues to alleviate eating disorders.

Social stressors can negatively impact puberty and sexual maturation in male and female humans and rodents (Bentefour and Bakker, 2024, 2024; Goldstein et al., 2010; Jaric et al., 2019). Stress exposure is associated with decreased levels of gonadal hormones and may be independent of stress hormone levels (i.e. cortisol) (Pletzer et al., 2021). Female hormonal functions are moderated by stress and influence stress-induced eating behaviors, revealing a possible molecular network mediating stress exposure and the subsequent eating response (Anversa et al., 2020; Calvez and Timofeeva, 2016; D’Souza and Sadananda, 2017; Goldstein et al., 2010; Jaric et al., 2019; Laham et al., 2022; Ma et al., 2020; Rocks et al., 2023). The interactions between chronic social isolation stress, gonadal hormones, naturalistic behaviors, and maladaptive coping have yet to be discovered.

The current study aimed to identify the longitudinal impact of social isolation stress on naturalistic behaviors in a rat model of stress-induced emotional hyperreactivity and binge-like eating to address these unresolved questions. Beginning with longitudinal monitoring of naturalistic behaviors in rats throughout adolescence, we identified several behaviors impacted by social isolation stress. Considering the relevance and efficacy of integrated behavioral z-scores, we observed social isolation stress increased emotional reactivity. After providing the rats with a Western-like high fat diet, we found that locomotor behaviors such as adolescent jumping and twitching were predictive later in life binge-like coping strategies. Understanding behavioral sensitivities to early life social isolation stressors not only helps improve predictive power but can unmask possible neurocircuitries contributing to later in life maladaptive emotional overeating and aids in establishing a foundation for future longitudinal preclinical investigations.

## METHODS

### Animals

All the experiments were performed following animal protocol 20-171, approved by the Institutional Animal Care and Use Committee (IACUC) at the Loma Linda University School of Medicine. This study follows the ARRIVE guidelines for reporting animal research (Percie Du Sert et al., 2020; Vega-Torres et al., 2022). When the rats were not in the home cage (see Home Cage Monitoring System below), they were kept in typical housing conditions (21 ± 2 °C, relative humidity of 45%, and a 12-hour light/dark cycle with lights on at 7:00 AM). All rat body weights, and food consumption were recorded weekly.

### Experimental Groups

Twenty-four female adolescent Lewis rats (postnatal day, PND, 15) were acquired from Charles River Laboratories (Portage, MI). Rats were habituated to housing conditions with 12-h light/dark cycles for at least one week before the initiation of the experiments. Females are underutilized in preclinical and translation research due to a common misconception about hormonal contribution to variability in rodent behaviors. Thus, leading to insufficient treatments for humans, and false assumptions about the underlying female physiology being negatively impacted by different disease states discussed in depth by Beery, 2018 and Shansky and Murphy, 2021. We wanted to address this health disparity by exclusively using females. Subjects were matched by body weight and weaned at PND 22. On the weaning day, rats were either pair-housed with two rats per cage (Paired, n=12) or single-housed (Isolated, n=12). This experimental group design is meant to reproduce early life stress induced by continuous social isolation stress. The rats were given *ad libitum* access to water and a purified diet (product no. F7463; Bio-Serv, Frenchtown, NJ). This diet was used to provide continuity between our studies.

### Longitudinal Study Design

We used home cage monitoring to assess various behaviors in the Paired and Isolated rats longitudinally. This portion of the study continued daily for seven weeks beginning at weaning (PND 22) until the rats transitioned into young adulthood (PND 72). The study timeline is summarized in **Figure 1**.

**Figure 1.**
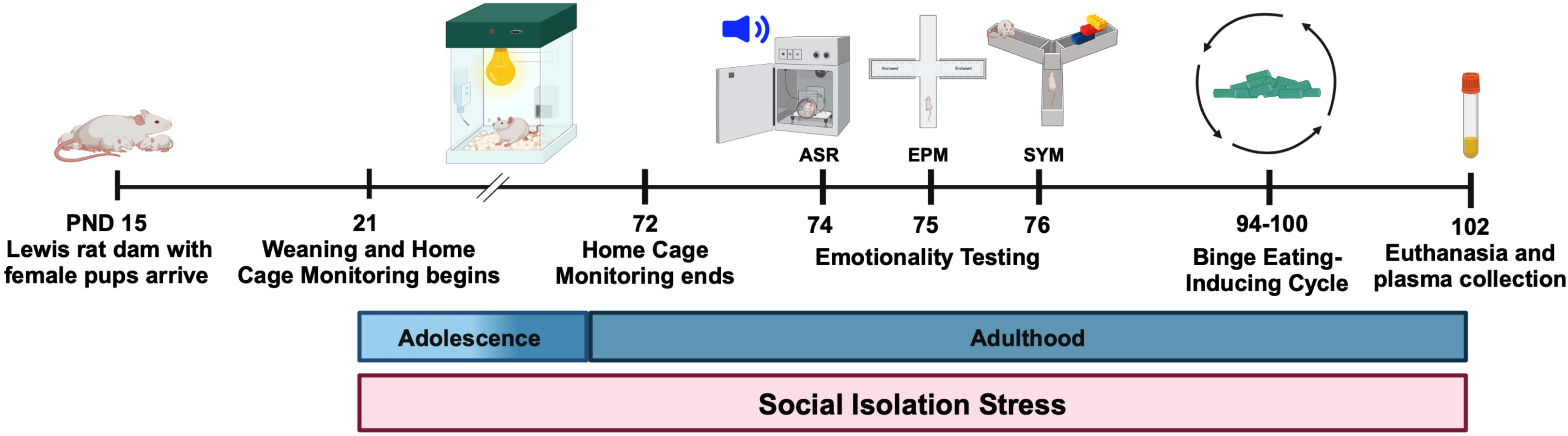
Experimental Timeline. Female Lewis rat pups acclimated for one week prior to weaning at PND 21. All animals were placed in Paired or Isolation housing. They were immediately placed into home cages for continuous overnight and weekly monitoring. After seven weeks, all animals underwent behavioral testing and subsequently engaged in our binge eating protocol.

### Home Cage Monitoring System

Noldus PhenoTyper cages were used for behavioral assessment instrumented observation (Noldus Information Technology BV, Wageningen, the Netherlands). The home cage monitoring system consists of a bottom plate that represents a black square arena; four replaceable transparent walls with ventilation holes at the top; an illuminated shelter that can be controlled with a hardware control module to switch the light automatically on when the rat enters the shelter, or choose a specific shelter entrance; a top unit that contains an infrared sensitive camera with three arrays of infrared light-emitting diode (LED) lights, and a range of sensors and stimuli, including adjustable light conditions to create a day/night cycle, the single tone for operant conditioning test, or the white spotlight for approach-avoidance behavior testing. Instrumented home cages allow testing of different behavioral characteristics of laboratory rodents in an environment like their home cage. Dark bedding (Purina Yesterday’s News Clumping Paper Cat Litter Unscented, Low Dust, Neenah, WI) was used to facilitate detection. The rats were observed in the home cages during the dark cycle for 12 h, 07:00 PM to 07:00 AM. The entire observation period was recorded and analyzed using EthoVision XT video tracking software, including the Rat Behavior Recognition module (Noldus Information Technology). EthoVision XT video tracking in conjunction with the Rat Behavior Recognition module enabled unbiased detection of digital biomarkers of social isolation stress effects on adolescent development. We measured grooming, jumping, supported rearing, unsupported rearing, twitching, sniffing, walking, resting, eating, and drinking cumulative duration. In addition, we measured eating frequency using feeding monitors incorporating a beam break device (Noldus). All independent naturalistic behaviors were examined hourly and weekly. To evaluate the independent behavior’s ability to predict adulthood emotionality and binge eating in adulthood, we pooled (Eraslan et al., 2023; Škop et al., 2023) each individual behavior’s data together to create a single value representing adolescence.

### Residual Avoidance

Adapted from Prevot et al. (2019), we used an aversive LED spotlight in the home cage to assess anxiety-like and food seeking behaviors. See Supplementary Figure 1 for the LED spotlight location and zone designations. We calculated residual avoidance (RA), i.e. the response to the aversive light challenge in the food zone (Eq. 1) and the shelter zone (Eq. 2), where “Time_[Time span],FZ_” and “Time_[Time span],SZ_” represent the total time spent in the food zone and shelter zone, respectively, in the specified time span. RA alludes to an rats’ response to dealing with a conflict.

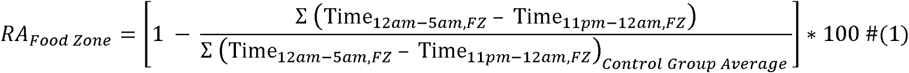

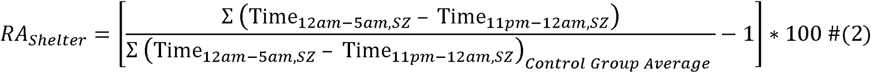

### Behavioral Assays

All behavioral assays were conducted during week eight beginning at PND 74 in the following order: acoustic startle reflex, elevated plus maze, and social Y maze. We have previously and consistently performed these behavioral tasks in this order to minimize carry over effects. Videos recordings were analyzed using Ethovision XT tracking software (Noldus Information Technology).

### Acoustic Startle Reflex (ASR)

The ASR experiments were performed using the SR-Lab acoustic chambers (San Diego Instruments, San Diego, CA, United States). ASR magnitudes were measured by placing rats in startle enclosures with sensors (piezoelectric transducers) that convert small movements to voltage. Thus, the magnitude of the change in voltage represents the size of the ASR. Acoustic stimuli intensities and response sensitivities were calibrated before starting the experiments. Our group has previously described the ASR protocol (Sharafeddin et al., 2024; Vega-Torres et al., 2018). Briefly, experimental sessions lasted 20 minutes and began with a 5-min habituation period (background noise = 60 dB). The rats were then presented with a series of 30 tones 105 dB) using a 30-s intertrial interval (ITI). The acoustic stimuli had a 20-ms duration. Subsequently, the rats were returned to their home cages. Enclosures were thoroughly cleaned and dried following each session. Averaged and maximum ASR magnitudes were normalized by weight to eliminate confounding factors associated with body weight (weight-corrected ASR = maximum startle magnitude in mV divided by body weight on testing day).

### Elevated Plus Maze (EPM)

The elevated plus maze (EPM) evaluates anxiety-like behaviors and captures a rodent’s instinctual desire to explore novel areas (Walf and Frye, 2007). Avoidance of the open arms is considered a measure of anxiety. The near-infrared (NIR)-backlit EPM apparatus (catalog #ENV-564A, MedAssociates) consisted of two opposite open arms (50.8 x 10.2 cm) and two enclosed arms (50.8 x 10.2 x 40.6 cm) that were elevated 72.4 cm above the floor. The junction area between the four arms measured 10 x 10 cm. A raised edge (0.5 cm) on the open arms provided additional grip for the rats. Behaviors were recorded in a completely dark room. This NIR maze is specifically designed for video-tracking applications and eliminates variables associated with shadow/lighting. At the beginning of each 5 min trial, the rat was placed on the central platform facing an open arm. The apparatus was thoroughly cleaned after each test session. Behaviors were monitored via a monochrome video camera equipped with a NIR filer fixed above the EPM. In addition to the standard measures, we included extra measures derived from ethological analysis of behavior. The time spent on both types of arms, the number of entries into both types of arms, and stretch attend postures (SAPs) were determined using EthoVision XT tracking software (Noldus Information Technology; RRID: SCR_000441, RRID: SCR_004074). **OA** denotes Open Arms while **CA** represents Closed Arms in the elevated plus maze.

### Social Y-Maze (SYM)

Sociability was assessed with a modified Y maze implementing a protocol adapted from (Vuillermot, 2017). The test was previously used to measure rodent social interactions. The Social Y-Maze (Conduct Science, Maze Engineers, Skokie, IL) is a plexiglass Y-maze with a triangular center (8 cm sides) and three identical arms (50 cm x 9 cm x 10 cm, length x width x height). One arm functions as the start arm, while the other two have rectangular wire mesh cages to hold the live conspecific or a ‘dummy object’. The protocol was conducted using one social interaction test trial without habituation training. The rat was placed into the start arm and allowed to explore the maze for 9-min freely. An unfamiliar conspecific (same age as the test rat) was put into one of the rectangular wire mesh cages. At the same time, a ‘dummy object’ consisting of multicolored LEGO® pieces (Billund, Denmark) was placed into the other wire mesh cage. The conspecifics and ‘dummy objects’ were counterbalanced across arms and treatment groups. The maze was cleaned with 70% ethanol and allowed to dry between trials. A camera was mounted directly above the maze to record each test session

### Estrus Cycle

To monitor estrus cyclicity, we stained and evaluated vaginal smears at PND 53-60. We used distilled water to flush the vaginal canal and then placed the water onto a glass slide. The slides were allowed to dry and then stained with crystal violet. To evaluate vaginal cytology, we imaged the stained slides with a Keyence BZ-X710 (Keyence Corporation, Osaka, Japan) brightfield microscope at 20X and 40X magnifications. We used vaginal cytology to stage each rat. Estrus = greater of cornified epithelial cells; Metestrus = greater levels of cornified epithelial cells and leukocytes; Diestrus = greater levels of leukocytes, low levels of nucleated epithelial cells, and absence of cornified epithelial cells; and Proestrus = greater levels of nucleated epithelial cells with low levels of cornified epithelial cells and absence of leukocytes by off (Ajayi and Akhigbe, 2020). Staging was determined by an experimenter blinded to the experimental conditions.

### Phenotypic Z-Scores

Z-scores were calculated by subtracting the mean of a behavioral measure from individual data points and dividing by the standard deviation (Eq. 3), where X is an individual value for an observed parameter, μ and σ denote the arithmetic mean and standard deviation of a user-defined baseline. This allows for the transformation of raw behavioral data into a standard normalized scale, facilitating meaningful comparisons between behavioral metrics and enhancing the interpretability of results (Meng et al., 2020; Sharafeddin et al., 2024).

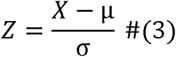

Individual z-score values for all 10 behaviors were averaged for each rat to produce an individual phenotypic score, and subsequently averaged to produce group phenotypic z-scores. The population mean and standard deviation were derived using the Paired group as baseline, allowing us to measure the deviation of Isolated phenotypic z-scores from the Paired group.

### Principal Component Analyses

The process consists of normalizing the data, generating principal components (PCs), and performing parallel analysis. Principal components were calculated using GraphPad Prism 10. Parallel analysis utilizes Monte Carlo simulations (1000 simulations; Percentile level of 95%) to select PCs with eigenvalues greater than the mean of corresponding PCs at the user-defined percentile level. Briefly, PCA was conducted on all ten behaviors for each week (Weeks 1-7) to visualize Paired versus Isolated clusters. Each week contained at least two PCs contributing to greater than 75% variability, which were used for subsequent analyses.

### Emotionality Z-Scores

Emotionality of rodents is best understood when integrating multiple behavioral outcomes into a single standardized score. Emotionality is a way of defining anxiety-and depressive-like states (Guilloux et al., 2011). Figure 4A depicts a graphical calculation pipeline for our emotionality z-score. We used z-normalized results from the complementary behavioral assays (ASR, EPM, and SYM) to determine the impact of social isolation stress on emotional domain. Z-scores were calculated from testing parameters measuring startle responses, sociability, avoidance, and ambulation. Higher z-scores are indicative of increased emotionality. We first classified the sub-readouts according to each behavioral test: ASR (mV__Max_, T-Max, and mV__Avg_); EPM (Cumulative Duration, Frequency, and Latency to the OA or CA, and SAP); and SYM (Cumulative Duration and Frequency with the Object or Conspecific, and SAP). Then, we performed Principal component analyses (PCA) on the behavioral test metrics to determine specific sub-readouts contributing to group variability. The three sub-readouts corresponding to the highest variation in PC1 were standardized and used to generate z-score averages. A comprehensive emotionality z-score was obtained by summing all metrics scores and dividing them by the number of total behavioral scores.

### Binge-like Eating

Following the behavioral testing, rats underwent a binge-eating disorder (BED)-like paradigm adapted from prior studies Czyzyk et al. (2010) and Sharafeddin et al. (2024). All rats were provided an ingredient-matched western-like diet (41% kcal from fat; WD) continuously for 48 hours. The initial exposure limits any novelty effect induced by exposure to a new diet. At the end of the 48-h period, the WD was replaced with the original purified diet for the remainder of the week. The following week, the diet manipulation was repeated, but rats were only exposed to WD for 24 hours. The 24-h period began at 8:00 AM until 8:00 AM the next day. Food consumption was measured 2.5-h and 24-h after the WD was provided. Isolated rats that ate significantly more than the Paired rats at 2.5 h were classified as binge-eaters.

### Euthanasia

All rats were anesthetized using isoflurane and injected intraperitoneally with Euthasol (150 mg/kg; Virbac, Fort Worth, TX). They were then perfused transcardially with ice-cold (4°C) phosphate buffered saline (PBS) using the Perfusion Two™ system (Leica Biosystems, Chicago, IL).

### Measuring corticosterone and gonadal hormone concentrations using ELISA

Blood was collected through cardiac puncture in EDTA–coated tubes. Blood samples were centrifuged at 1000 rpm for plasma fraction collection and stored at -80°C until further testing. Measurement of circulating hormone concentrations in plasma were measured using ELISA kits purchased from Enzo Life Sciences (Farmingdale, NY, USA) according to manufacturer’s instructions, which included: corticosterone (Cat. No. ADI-900-097), estrogen (17β-Estradiol) (Cat. No. ADI-900-174), progesterone (Cat. No. ADI-900-011), and luteinizing hormone (LH) (Cat. No. ENZ-KIT107). Our group previously used ELISA for circulating hormones and molecules (Kalyan-Masih et al., 2016). Corticosterone and progesterone samples were treated with a steroid displacement reagent (included in kit) to free unbound steroid in samples containing steroid-binding proteins (i.e. plasma). Samples were diluted 1:10 in assay buffer (1:30 for corticosterone). Plate absorbances were measured at 405 nm with 570 nm reference wavelength (450 nm and 570 nm reference for LH) using the SpectraMax i3X detection platform (Molecular Devices, Sunnyvale, CA). Concentrations were obtained using 4PLC interpolation from the standard curve and corrected by the dilution factor.

### Statistical Methods

Statistical analyses were performed using GraphPad Prism 10. Two-way analysis of variance (ANOVA) was used, when appropriate, to examine the effect of social isolation stress, time, and interaction between factors on outcome measures. Multiple comparisons were made using Tukey’s test. The Shapiro-Wilk and Brown-Forsyth tests were used to assess the assumptions of normality and homoscedasticity, respectively. The ROUT method was used to investigate outliers. Differences were considered significant for *p*<0.05. The data is shown as the mean ± standard error of the mean (S.E.M.) unless otherwise stated. Spearman correlations were run on integrated/composite data and Pearson correlations on non-integrated data.

## RESULTS

### Longitudinal composite phenotypic z-scores of naturalistic behaviors reveal social isolation stress-induced shifts in adolescent behavioral developmental trajectories

To comprehensively assess the longitudinal impact of social isolation stress on naturalistic behaviors, we continuously tracked single and paired housed female adolescent rats and standardized all 10 behaviors to calculate weekly composite phenotypic z-scores. We found that social isolation stress deviated developmental trajectories and that several naturalistic behaviors were more susceptible to social isolation stress. Weekly composite phenotypic z-scores (Figure 2B, S. Table 1) revealed social isolation stress-induced significant deviations from the pair-housed controls stress effect (*F* (1,22) = 15.91, *p* = 0.0006), time effect (*F* (3,064) = 48.97, *p* < 0.0001), and interaction effect (*F* (6,124) = 5.388, *p* < 0.0001). To exemplify the increase in phenotypic profiles of the isolated rats, we pooled the phenotypic z-score data into a singular i.e. integrated, score representing the entirety of adolescence (*t* (22) = 3.218, *p* = 0.0040) (Figure 2C). PCA unsupervised clustering of all 10 home cage behaviors for each week showed a distinct separation between Paired and Isolated rats (Supplementary Figure 2). To provide a quantitative measure of these differences, we compared PC1 and PC2 scores between Isolated and Paired rats for each week. We found that Isolated rats had altered PC1 (Supplementary Figure 3, S. Table 2) and PC2 scores (Supplementary Figure 4, S. Table 2) suggesting the behavioral profiles corresponding to the Isolated rats are distinct from the Paired.

**Figure 2.**
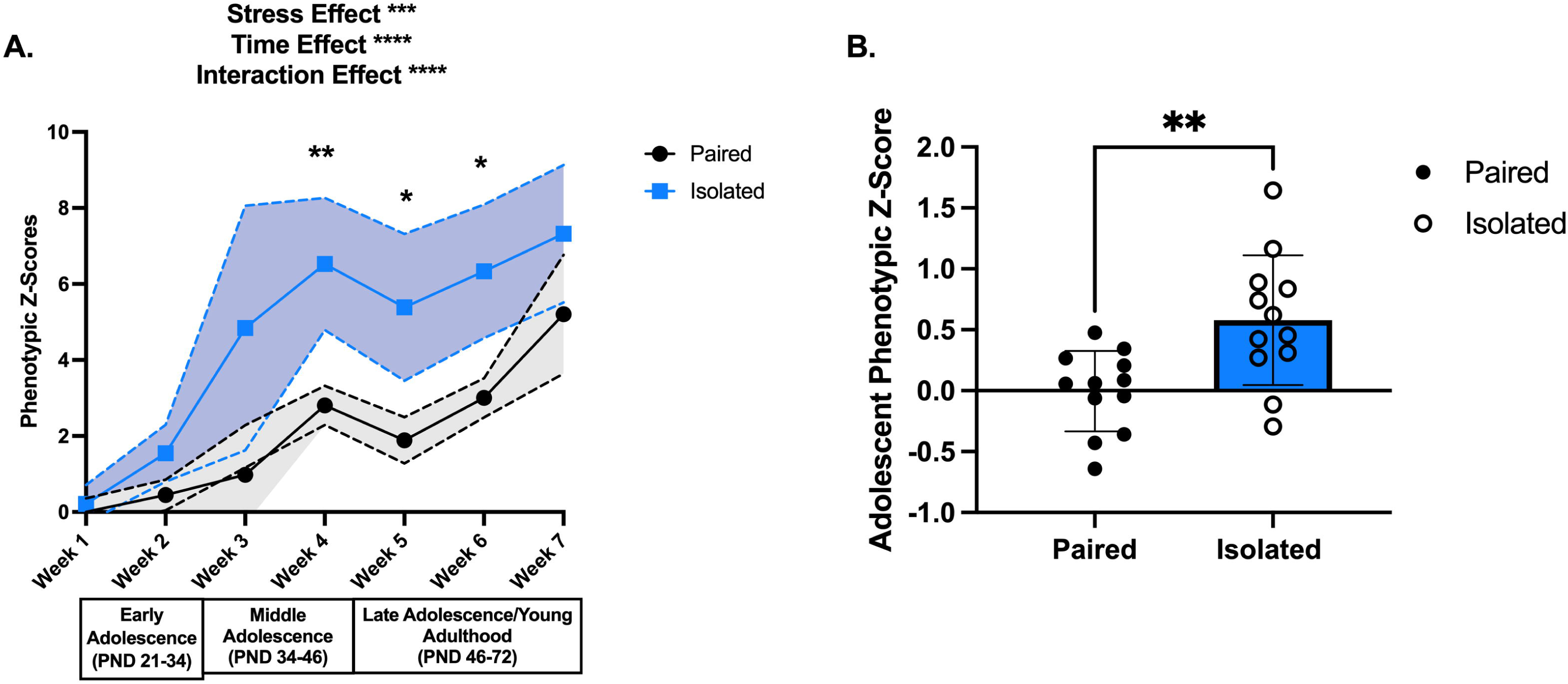
Isolation housing induces greater shifts in phenotypic z scores throughout adolescence compared to paired housing. **(A)** Shows the phenotypic z scores for each week for the Paired (black) and Isolated (blue) animals using Paired week 1 as the population baseline. Dashed lines and shaded areas reflect 95% confidence intervals. **(B)** Adolescent phenotypic z scores calculated from all the behaviors demonstrated statistically significant increase in the Isolated compared to the Paired animals. Paired and Isolated, n=12. Asterisks (*) signify statistically significant differences when comparing Paired and Isolated animals for that week. Data presented with SEM unless otherwise stated. * p < 0.05, ** p < 0.01, *** p < 0.001, **** p <0.0001; Mixed Effects ANOVA, Sidak Post hoc comparisons. See Table 1 for detailed statistics.

In addition to evaluating composite and integrated phenotypic profiles, we examined which behaviors were susceptible to isolation stress. Jumping, unsupported rearing, cumulative sniffing duration, and twitching frequency were significantly increased in isolated rats compared to paired ones (Supplementary Figure 5, S. Table 3). These behaviors are therefore considered more sensitive to the experimental manipulations and reflect heightened susceptibility to social isolation stress. To determine the contribution of these behaviors to the behavioral profile differences between the Isolated and Paired groups, we recalculated phenotypic z-scores with either the significantly shifted behaviors (*t* (22) = 4.848, *p* < 0.0001) (Supplementary Figure 6A) or the remaining non-impacted behaviors (*t* (22) = 0.7506, *p* = 0.4608) (Supplementary Figure 6B). These results demonstrate that the subset of significantly impacted behaviors is sufficient to capture overall isolation induced-behavioral profiles by maintaining inter-group differences. To examine whether these differences were due to asynchronized estrus cyclicity, we used vaginal cytology during the transition to puberty (PND53-60) and found no significant differences (Supplementary Figure 7). Altogether, our unbiased methodology revealed longitudinal changes in the behavioral profiles of socially isolated adolescent rats.

### Socially isolated rats endure an aversive spotlight zone in favor of the food zone

To probe food seeking and anxiety-like behaviors within the home cage, we adapted a paradigm of residual avoidance where an aversive light is turned on during the dark cycle illuminating the center of the cage. We found that social isolation stress led to a persistent preference for the food zone despite the presence of an aversive stimulus. Supplementary Figure 8 and S. Table 4 displays the changes in home cage behaviors before, during, and after the spotlight hour from all seven weeks. The isolated rats spent a comparable amount of time in the shelter zone (*F* (1,141) = 2.283, *p* = 0.1330) (Figure 3A), but displayed a significant preference for the food zone when the spotlight challenge terminated (*F* (1,141) = 4.014, *p* = 0.0470) (Figure 3B) and during the spotlight hour (*F* (1,141) = 7.545, *p* = 0.0068) (Figure 3C). See S. Table 5 for detailed statistics. Taken together, these data suggest social isolation stress leads to preference of the food zone during and even after the threat has been terminated, indicating a possible coping strategy. Appreciating the home cage’s ability to yield longitudinal naturalistic behavioral and anxiety-like deficits in our rats, we aimed to compare these alterations with a conventional battery of behavioral tests related to emotionality.

**Figure 3.**
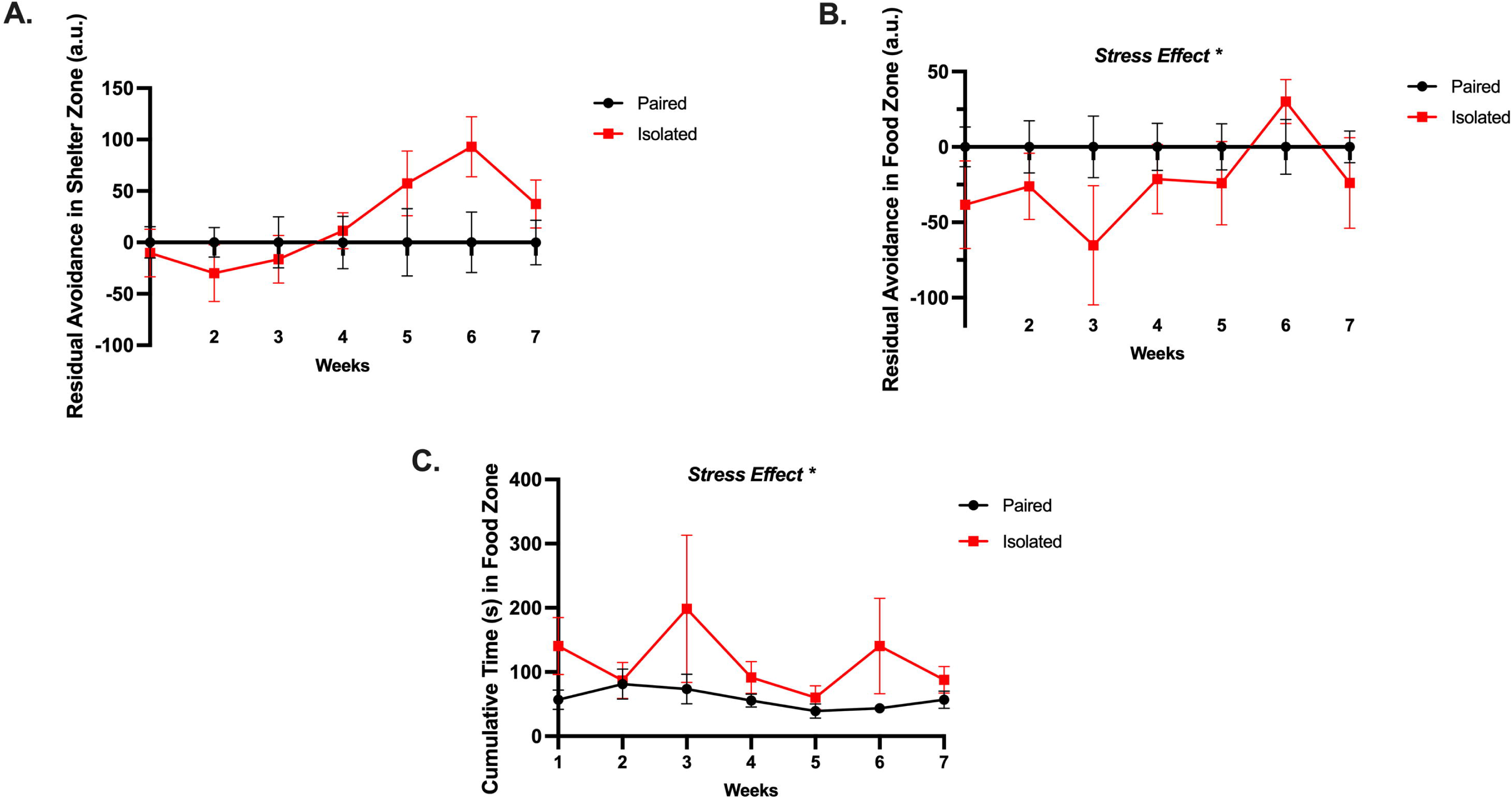
Isolated animals endure as aversive spotlight area in favor of the food zone with no differences in shelter zone preference. **(A)** Isolated and Paired animals do not show any differential preferences for the shelter zone once the spotlight has been turned off. **(B)** Isolated animals endure the spotlight zone in favor of the food zone after when the spotlight is discontinued. **(C)** Isolated animals spend more time in the food zone when the spotlight is turned on. See Residual Avoidance Formula in Materials and Methods section. Paired and Isolated, n=12. Data presented with SEM. Asterisks (*) signify statistically significant differences when comparing Paired and Isolated animals for that week. * p < 0.05, ** p < 0.01, *** p < 0.001, **** p <0.0001; TWO-WAY ANOVA, Sidak Post hoc comparisons.

### Socially isolated rats display elevated emotionality

To evaluate emotionality, we integrated multiple behavioral readouts (Supplemental Figures 9, 10, and 11; S. Table 6, 7, 8, and 9) from traditional independent behavioral assays to calculate an integrated emotionality z-score (Guilloux et al., 2011). Averaged z-scores revealed a significant increase in ASR (*t* (22) = 3.693, *p* = 0.0013) (Figure 4A) and SYM (*t* (20) = 2.139, *p* = 0.0449) (Figure 4B), but not EPM (*t* (22) = 1.008, *p* = 0.3243) (Figures 4C), in isolated compared to paired rats. To calculate an integrated emotionality z-score, we averaged the individual behavioral z-scores demonstrating an increase in emotionality of socially isolated rats (*t* (22) = 4.017, *p* = 0.0006) (Figure 4D, Supplementary Figure 12). To address the lack of knowledge connecting naturalistic behaviors with emotionality and affective valence, we correlated each shifted behavior with the emotionality z-score and discovered only adolescent jumping cumulative duration to be associated with emotionality (*Spearman r* = 0.4922, *p* = 0.0146) (Figure 4E). Considering the association between heightened emotionality and maladaptive coping strategies such as binge eating (Bodell et al., 2019; Müller et al., 2014; Sanzari et al., 2023; Sulkowski et al., 2011), we evaluated the rats’ responses to a Western-like high fat diet (WD).

**Figure 4.**
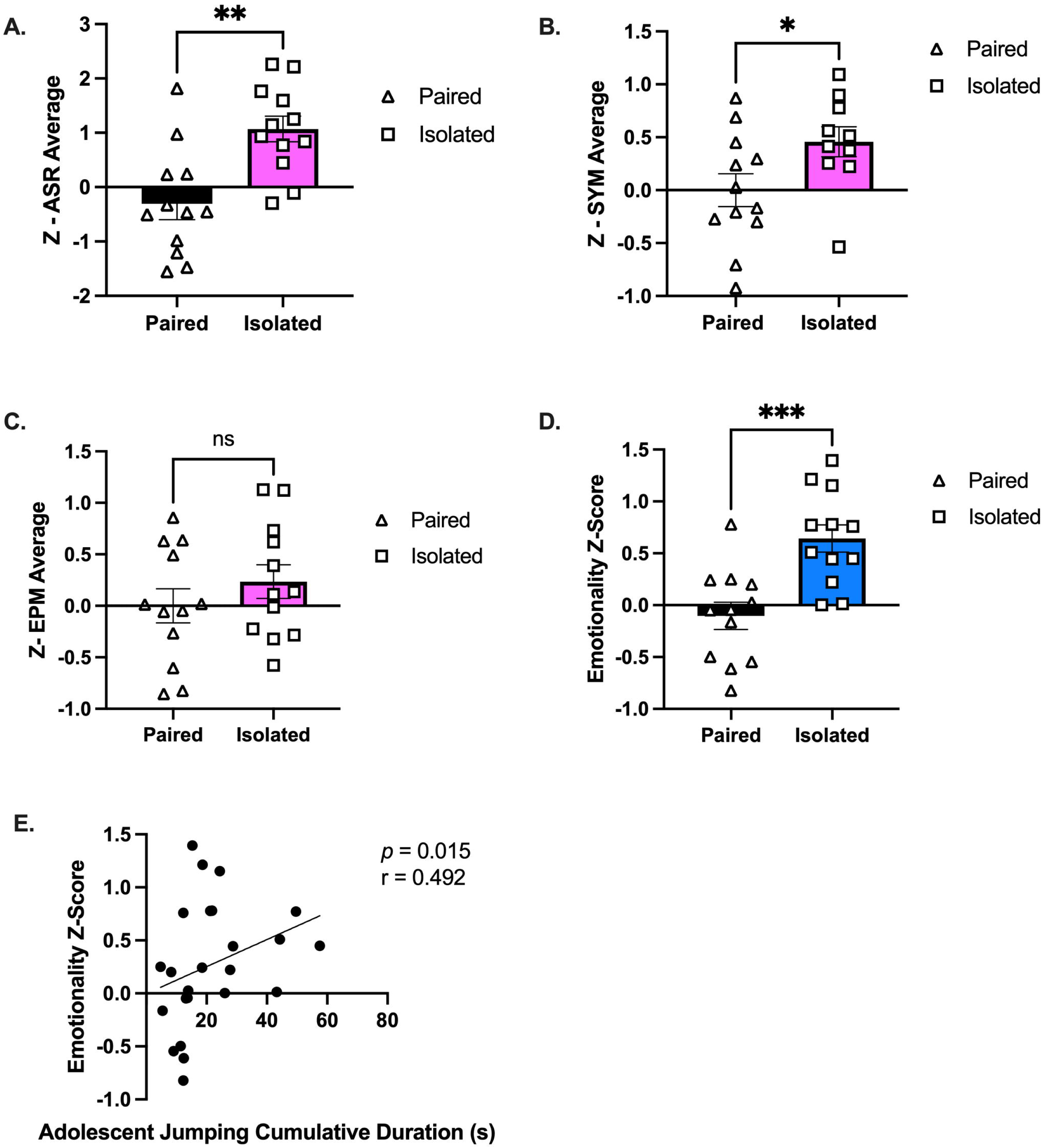
Isolated animals display an increased emotionality composite z-score compared to Paired animals. **(A-C)** Average of standardized z-scores for ASR, EPM, and SYM, respectively. **(D)** Overall emotionality z score. **(G)** Emotionality z score is correlated with adolescent jumping behaviors when analyzed with all the animals. Paired and Isolated, n=12. Data presented with SEM. * p < 0.05, ** p < 0.01, *** p < 0.001, **** p <0.0001; Student t-and Mann-Whitney tests were used for parametric and nonparametric data, respectively. Spearman Correlations were used for emotionality z score correlation analyses. p and r values are presented.

### Naturalistic behaviors predict Western diet food consumption and binge-like eating behaviors emerge in a subgroup of isolated rats

We implemented a paradigm to induce binging without food restriction, consisting of 24-h continuous access to the WD beginning in the morning (light cycle) until the following morning. We identified adolescent naturalistic behaviors predictive of binge-like eating. Initial experiments using this binge-like paradigm revealed more social isolation stress-induced bingeing in females compared to males (Supplementary Figure 13, S. Table 10). Interestingly, adolescent twitching frequency (*Pearson r* = 0.5552, *p* = 0.0049) (Figure 5A) and adolescent jumping cumulative duration (*Pearson r* = 0.5140, *p* = 0.102) (Figure 5B) were positively correlated with WD consumption during the initial binge-like eating session (2.5-h period). To tease out the effect of social isolation stress, correlation analyses were conducted on each group separately. We found that the association between adolescent twitching and WD consumption was maintained in the isolated rats (*Pearson r* = 0.657, *p* = 0.020), but was lost in the paired group (*Pearson r* = 0.405, *p* = 0.265) (Figure 5C), demonstrating that social isolation stress is driving the relationship between these behaviors. We observed a similar trend with jumping cumulative duration, although it did not reach statistical significance (Isolated, *Pearson r* = 0.570, *p* = 0.053; Paired, *Pearson r* = 0.047, *p* = 0.884) (Figure 5D).

**Figure 5.**
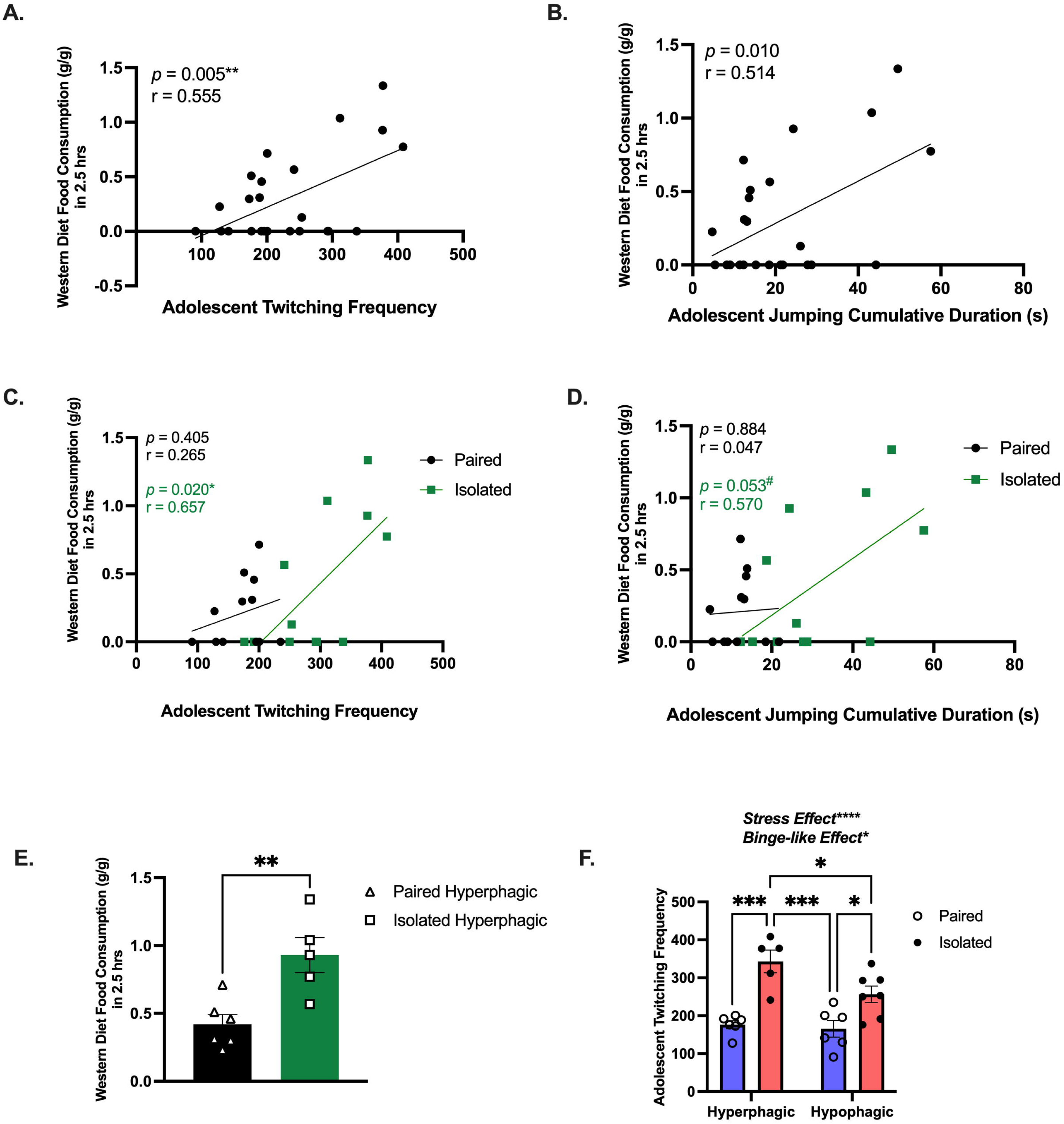
Isolation exacerbates bingeing and is predicted by twitching behavior. **(A)** Correlation using all animals between Adolescent twitching frequency and the 2.5hr WD consumption measurement. (n=24) **(B)** Correlation between Adolescent jumping cumulative duration and the 2.5hr WD consumption measurement. (n=24). **(C)** Correlation between Adolescent twitching frequency and the 2.5hr WD consumption measurement for Isolated (n=12) and Paired (n=12). **(D)** Correlation between Adolescent jumping cumulative duration and the 2.5hr WD consumption measurement for Isolated (n=12) and Paired (n=12). **(E)** These data demonstrate subpopulations of animals more likely to consume the WD. **(F)** Adolescent twitching frequency can distinguish high and low eaters (n=5-6) for each group and paired and isolated animals. (n=12) for each group. Data presented with SEM. ^#^ is approaching significance, * p < 0.05, ** p < 0.01, *** p < 0.001, **** p <0.0001; Student t-and Mann-Whitney tests were used for parametric and nonparametric data, respectively. Pearson Correlations were used when comparing naturalistic behaviors with food consumption. p and r values are presented. TWO-WAY ANOVA, Sidak Post hoc comparisons. We called the statistically significant increase between High *and Low eaters a Binge-like Effect*.

Next, we investigated behavioral shifts based on food intake during the binge-like eating paradigm. Rats that consumed above their respective group average were reclassified as ‘hyperphagic’. In doing so, we found that isolated hyperphagic rats consumed significantly more than the paired hyperphagic rats during the 2.5-h binge-eating period (*t* (9) = 3.603, *p* = 0.0057) (Figure 5E). To determine whether adolescent twitching, jumping, or emotionality could uniquely distinguish between the hyper-and hypo-phagic rats, we evaluated each domain. Notably, only adolescent twitching frequency was significantly elevated in the hyperphagic rats compared to the hypophagic rats (*F* (1,20) = 5.032, *p* = 0.0364) and stress-effect (*F* (1,20) = 35.28, *p* < 0.0001) (Figure 5F), indicating that adolescent twitching frequency could be predictive of, not only stress-induced bingeing, but possible bingeing in general. See S. Table 10 for detailed statistics. Collectively, our model uncovered subpopulations of rats that are more vulnerable to binge-like eating when re-introduced to a WD and that social isolation stress exacerbates this vulnerability. These behavioral differences beg the question of the possible molecular players underlying these observations.

### Corticosterone levels are reduced in socially isolated rats

We measured circulating levels of corticosterone, estrogen, progesterone, and luteinizing hormones. We determined that social isolation stress promoted a reduction in corticosterone and that changes in hormone levels are associated with adolescent naturalistic behaviors. Figure 6A shows that circulating levels of corticosterone were significantly lower in isolated rats compared to the paired group (*t* (20) = 4.006, *p* = 0.0007) with no differences in gonadal hormones (Supplementary Figure 14 A-C). PCA further revealed indiscriminate differences between the isolated and paired rats when using the gonadal hormones as variables (Supplementary Figure 14D). Unsupported rearing cumulative duration was positively correlated with estrogen (Supplementary Figure 15A) and progesterone (Supplementary Figure 15B). Progesterone was also positively correlated with corticosterone (Supplementary Figure 15C). Curiously, when delineating these effects from paired and isolated rats, the association between unsupported rearing and estrogen was sustained only sustained in the paired rats (Supplementary Figure 15D), while the association between unsupported rearing and progesterone was observed only in the isolated rats (Supplementary Figure 15E). Corticosterone levels were negatively correlated with the emotionality z-score (*Spearman r* = -0.4179, *p* = 0.0421) (Figure 6B), adolescent shifted behaviors score (*Spearman r* = -0.428, *p* = 0.042) (Figure 6C), adolescent twitching frequency (*Pearson r* = -0.4627, *p* = 0.0228) (Figure 6D), and adolescent jumping cumulative duration (*Pearson r* = -0.4038, *p* = 0.0503) (Figure 6E). See S. Table 12 for overview of statistics. Together, these data confirm the influence of corticosterone on emotionality, highlighting the advantage of using longitudinal home cage observations to identify biobehavioral markers. Altogether, our model successfully recapitulated aspects of critical molecular imbalances found in eating disorders.

**Figure 6.**
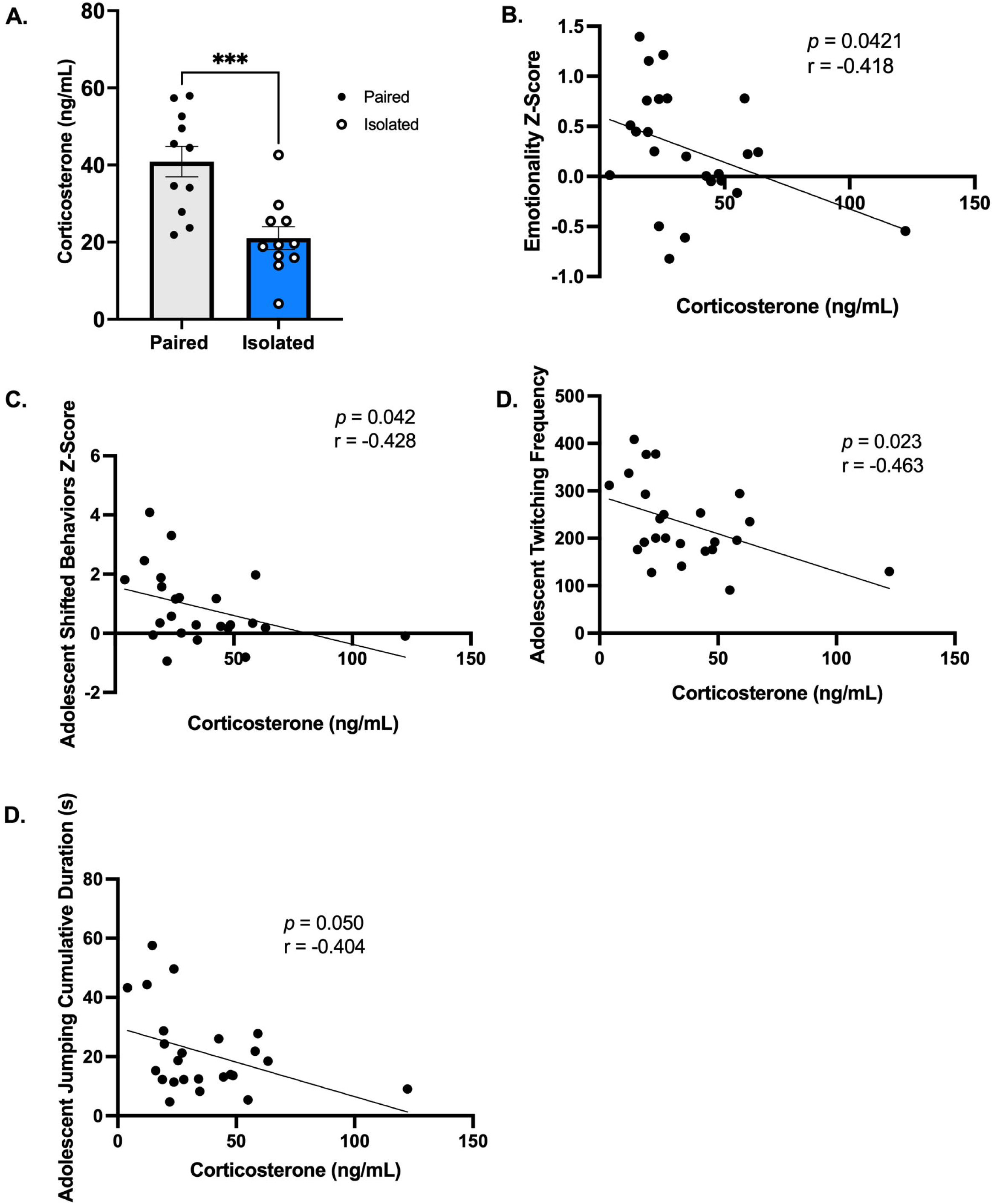
Corticosterone is blunted in Isolated animals and is correlated behavior and emotionality. **(A)** Isolation housing blunts corticosterone levels (n=11, Paired and Isolated). **(B-E)** Corticosterone levels are correlated with emotionality z scores, adolescent shifted behaviors, twitching frequency, and jumping cumulative duration. Data presented with SEM. * p < 0.05, ** p < 0.01, *** p < 0.001, **** p <0.0001; Student t-and Mann-Whitney tests were used for parametric and nonparametric data, respectively. Pearson Correlations were used when comparing naturalistic behaviors and Spearman for Emotionality z scores. p and r values are presented.

## DISCUSSION

The present study determined (1) the feasibility of longitudinal monitoring of unbiased naturalistic behaviors in rats, (2) the value of using standardized composite z-scores to compare phenotypic profiles of rats exposed to different environmental manipulations through time, (3) the effectiveness of an aversive spotlight to identify anxiety-like avoidant behaviors, (4) changes in emotional reactivity in response to adolescent social isolation stress, (5) the use of naturalistic home cage behaviors to predict stress-induced binge-like eating, and (6) associations between behavior and circulating neurohormone levels.

Measuring rat behavior longitudinally offers several advantages that can contribute to identifying early risk biomarkers for stress-induced alterations. Longitudinal studies allow tracking behavioral changes over time to discover phenotypic variations and individual trajectories, allowing for a more nuanced understanding of how certain behaviors may be linked to brain pathologies or maladaptive coping strategies (Bodell et al., 2019; Febo et al., 2020; Klatzkin et al., 2024; Lacroix et al., 2023; Moorhead et al., 2003). Similarly, automated behavioral recognition software has many advantages over conventional behavioral testing, such as automatic detection and tracking of a diverse repertoire of rodent behaviors, reducing experimenter bias, minimizing human error, and enhancing reproducibility (Klibaite et al., 2022; Lee et al., 2020; Philip et al., 2010; Reinert et al., 2019). The automated recognition software utlized here enabled us to tease our specific behaviors more susceptible to chronic social isolation stress during adolescence. Using these approaches, we observed an increase in involuntary twitching, characterized by sudden muscle movements. Additionally, isolated rodents exhibited repeated, often frantic jumping. Persistent and exaggerated sniffing behavior was also noted, along with unsupported rearing, where the rodents frequently stood on their hind legs without external support. These findings highlight the considerable strain that social isolation stress places on the typical behavioral patterns of female rats. In addition to understanding specific ethological behavioral deficits induced by adolescent psychosocial stress, there is a critical need to develop methods that more appropriately capture endophenotypes. Composite scores are a translationally relevant measure of behavioral phenotypes and shifts in behavioral profiles (Guilloux et al., 2011). We have previously used home cages and phenotypic z-scores to investigate chronic stress effects in Lewis rats (Sharafeddin et al., 2024). Harnessing phenotypic z-scores, our study found that social isolation stress in female rats throughout adolescence shifted the developmental trajectories of naturalistic/home cage behaviors specifically by increasing these behaviors. There was a clear progressive deviation of the isolated phenotypic z-score from the paired group leading up to week 7, where they converge. It is possible the socially isolated females were no longer coping with isolation by excessively engaging in these behaviors. Rats transition to young adulthood at week 6-7 and this maturational advancement could have influenced behavioral outcomes. Despite this convergence, we know the overall negative impact of social isolation stress was maintained into young adulthood because of the evident differences in subsequent measures of emotionality, binging, and hormone levels. However, the precise reason for this convergence, whether biological or technical, remains unknown. Subsequent studies with more controls are needed to address these questions. Evidence supports the long-term detrimental impact of early-life social isolation in adult animals but neglects details about early behavioral deviations. It is reasonable that more behaviors were impacted but failed to be detected by the software. As behavior recognition software evolve, they will become more accurate and able to detect more behaviors (Van Dam et al., 2023). In summary, this approach can identify specific ethological behavioral patterns or changes that precede or predict subsequent neurodegenerative and neuropsychiatric alterations. Understanding these trajectories through time can aid with establishing predictive models to impede the progression of psychopathology while providing a behavioral profile indicative of the underlying neural landscape.

Furthermore, the aversive spotlight revealed that socially isolated rats tended to disregard the spotlight zone in favor of the food zone more than pair-housed rats, suggesting elevated food seeking behavior that maybe driven by enhanced anxiety and emotionality. Foundational studies employing the aversive spotlight to assess avoidant behaviors in mice demonstrated psychosocial stress led to consistent residual avoidance of the aversive spotlight area even after it had been turned off (Prevot et al., 2019). There is also evidence that stressed rodents will seek food in the presence of an aversive stimulus (Calvez and Timofeeva, 2016; Oswald et al., 2011). The current study is the first to our knowledge that has measured similar metrics in rats. The evident ethological and anxiety-like deficits induced by chronic social isolation stress was strengthened by emotionality testing.

Next, composite emotionality scores supported the phenotypic z-score findings that social isolation stress negatively impacts adolescent female rats. Independent assays revealed that socially isolated rats demonstrated elevated acoustic startle reactivity, anxiety-like behaviors, and reduced sociability consistent with our groups past findings (Kalyan-Masih et al., 2016; Sharafeddin et al., 2024). We calculated a composite emotionality score by aggregating behavioral metrics contributing most to variation in the data. This emotionality score suggests that socially isolated rats exhibit higher emotional reactivity compared to paired rats. Individuals with anxiety and depressive disorders, PTSD, and early life adversities commonly display alterations in emotional reactivity (Badour and Feldner, 2013; Lee-Winn et al., 2016). Furthermore, emotional reactivity could be a predictor of substance abuse, overeating, and other coping strategies (Ljubičić et al., 2023).

Binge eating is a highly prevalent form of maladaptive coping where an individual consumes a large amount of palatable food in a short period of time, accompanied by a sense of loss of control (*Diagnostic and statistical manual of mental disorders*, 2013; Hudson et al., 2007). Binge eating behaviors are associated with social stress exposure in humans and rodents (Klatzkin et al., 2024; Makri et al., 2022; Quilliot et al., 2019; Shin and Miller, 2012; Sulkowski et al., 2011; Wattick et al., 2023) and became particularly evident in adolescents who experienced social quarantine during the COVID-19 pandemic and were reported to have higher levels of emotional eating and had an increased risk of developing eating disorders (Hansen et al., 2021; Raventós et al., 2023; Trafford et al., 2023). Rat models are especially helpful in this context, as they allow for controlled manipulation of environmental and genetic factors to explore the underlying biological mechanisms of binge eating. The association between stress and binge eating is clear, but behavioral patterns predictive of disordered eating development and variation in individual responsivity to stress remain unclear. Velázquez-Sánchez et al. 2014 found that adult male rats with high trait motor impulsivity predicted the development of food addiction and binge-like behaviors when food was provided during the rodents’ dark cycle. However, traditional behavioral assays are aimed at testing a specific behavior or trait. Therefore, we used naturally motivated behaviors to characterize markers of over-eating susceptible phenotypes amplified by early life social isolation stress. Adolescent twitching frequency was the most salient behavioral predictor of binge-like eating. Trait and stress-induced twitching and jumping could be detected in their home cages without pretraining or habituation to the testing environments unlike other studies attempting to predict binge-eating. Traditional behavioral assays heavily restrict the range of alternative actions of the experimental animals to specifically test a behavior or decision. Others have suggested that investigators might miss or underreport stress-related repetitive behaviors in rodents because these behaviors often happen at night and stop when the experimenter enters the room (Krohn et al., 1999). Therefore, more detailed and reliable methods are needed to study abnormal behaviors in nocturnal animals. These limitations emphasize the utility of long-term observation cages to detect naturally motivated behaviors such as motor impulse without being subjected to predetermined choices. Altogether, our study and others reinforce the importance of both longitudinal and predictive studies to explain the individual response variability.

Specific ethological behaviors were positively associated with WD consumption, which was exacerbated in socially isolated rats. Adolescent jumping and twitching showed a stronger association with WD consumption than other shifted behaviors. Motor behaviors are energy-intensive behaviors which necessitates higher caloric intake to compensate for increased energy demand (Škop et al., 2023). This could explain the increased caloric consumption observed in our isolated rats/hyperphagic, which performed these behaviors considerably more than the paired. Studies in rodents have demonstrated social isolation stress-induced increased locomotion (Deal et al., 2021; Sullens et al., 2021) and that locomotion could be a predictor of later in life palatable food and ethanol consumption (Hsu et al., 2010; Karatayev et al., 2010; Morganstern et al., 2010; Rodríguez-Rangel et al., 2024). Twitching is a rapid and fast movement of the body, categorized as stereotypic (abnormal repetitive behavioral patterns) movement, which is thought to be adversely influenced by stress and theorized to be a self-alleviation strategy in suboptimal environments (*Recognition and Alleviation of Distress in Laboratory Animals*, 2008). While there is no consensus on increases or decreases in stereotypies across species, it is generally accepted that shifts in stereotypies are indicative of emotional state (Garner and Mason, 2002; Novak et al., 2016b; Turner and Bayne, 2023). Others have used home cages to observe how chronic stress can increase stereotypies, such as bar-mouthing in mice (Novak et al., 2016b, 2016a); these studies have short durations, lack sophisticated detection methods, and omit integrated behavioral analyses. Stereotypies, such as twitching, may arise due to dopamine dysregulation, as dopaminergic neurocircuits modulate activity, or movement behaviors (Belskaya et al., 2024; Chartoff et al., 2001; Gainetdinov et al., 2001; Garner and Mason, 2002; Leo et al., 2018). Social isolation stress can increase dopamine uptake in the striatum (Yorgason et al., 2016). Future studies on the role of neurotransmission dysregulation in mediating hyper repetitive movements will generate more therapeutic agents to ease the burden of psychosocial stress-induced binge eating. Unsurprisingly, there was large variability in the total amount of food consumed, regardless of housing conditions. The mechanisms underlying individuals’ variable response to social stress with food is unknown, therefore understanding the molecular mechanisms underlying ethological behaviors shifted by stress will supply prospective neural circuits to explore.

Measures of estrous cyclicity by vaginal cytology and circulating hormone levels did not yield an association with disordered eating or emotionality or display differences between groups. However, circulating corticosterone levels were reduced by the social isolation stress which could correspond to dysregulated HPA responsivity and has been seen in other social stress models (Gądek-Michalska et al., 2017). HPA responsivity can be reduced with repeated stress and may be rooted in hippocampal abnormalities (Algamal et al., 2018). Fluctuations in cortisol levels have been observed in women with stress disorders (Meewisse et al., 2007), and influence binge eating (Gluck et al., 2004). Blunted plasma corticosterone has also been observed in binge-like eating prone female rats (Calvez and Timofeeva, 2016). Stress disorders and stress related behaviors are linked with low circulating corticosterone levels in rats (Monari et al., 2024; Whitaker and Gilpin, 2015). This response may be unique to Lewis rats since they are known to have blunted HPA response compared to other rat strains (Cohen et al., 2006; Michaud et al., 2003). This study used female Lewis rats, which are highly susceptible to psychosocial stressors, display enhanced impulsivity and anxiety-like behaviors (Hamilton et al., 2014; Meyza et al., 2011; Sharafeddin et al., 2024). Our group will continue using this model to investigate stress susceptible phenotypes.

We acknowledge some limitations to the study design. Our primary psychosocial stressor was chronic social isolation by single housing our rats beginning at weaning (PND 21). However, subjects were separated from dams and placed into home cages without an acclimation period, which could have introduced an additional stressor. By measuring the deviation from the Paired group rather than from a separate control, the effect of the acute stressor imposed on the rats should be mitigated since both groups endured similar weaning conditions. Future studies should investigate the effects of weaning on behavioral phenotypes. This presents an additional stressor that could have influenced the rats’ developmental trajectories (Prevot et al., 2019). Additionally, while both groups were body weight-matched to ensure initial weights did not influence future eating outcomes, we did not ASR-match them as our previous studies (Kalyan-Masih et al., 2016; Ontiveros-Ángel et al., 2024; Sharafeddin et al., 2024; Vega-Torres et al., 2022, 2020, 2018). ASR matching could have introduced as additional acute stressor. These additional stressors could exacerbate the effects of social isolation or inadvertently confer overall stress resilience which is seen in rats and mice (Chaby et al., 2020; Cotella et al., 2020; Santarelli et al., 2017, 2014).

In summary, measuring rat behavior longitudinally offers advantages in understanding the developmental progression of behavior, identifying individual differences and trajectories, and detecting early risk markers predictive of coping strategies such as binge eating. Using this approach, we evaluated shifts in naturalistic behaviors in female rats exposed to continuous social isolation stress and showed how these dynamics help predict adult behaviors, unveiling a prospective avenue for improvements in personalized preventative and therapeutic interventions in the future. In doing so, we gained valuable insights into the temporal dynamics of naturalistic behaviors and their relationship to negative behaviors later in life. This study provides compelling evidence that composite phenotypic z-scores encompassing 10 naturalistic behaviors longitudinally offer valuable insights into organism-level behavioral profiling of chronic social isolation stress in rats and demonstrates the associations between ethological behaviors (Figure 7, Supplementary Figure 16). Phenomics approaches implementing high throughout phenotyping devices will garner more translationally relevant data from preclinical models. We successfully integrated multiple behavioral modalities to produce composite scores capable of wholistically evaluating differences in emotionality and phenotypic behavioral profiles within and between our rats. Finally, we added valuable insight into the interaction between circulating stress hormone and maladaptive coping strategies induced by continuous social isolation stress. Our results pave the way for utilizing integrated longitudinal observations in rodents to predict coping behaviors, such as binge-like eating, providing insight into the long-term effects of early life social stress.

**Figure 7.**
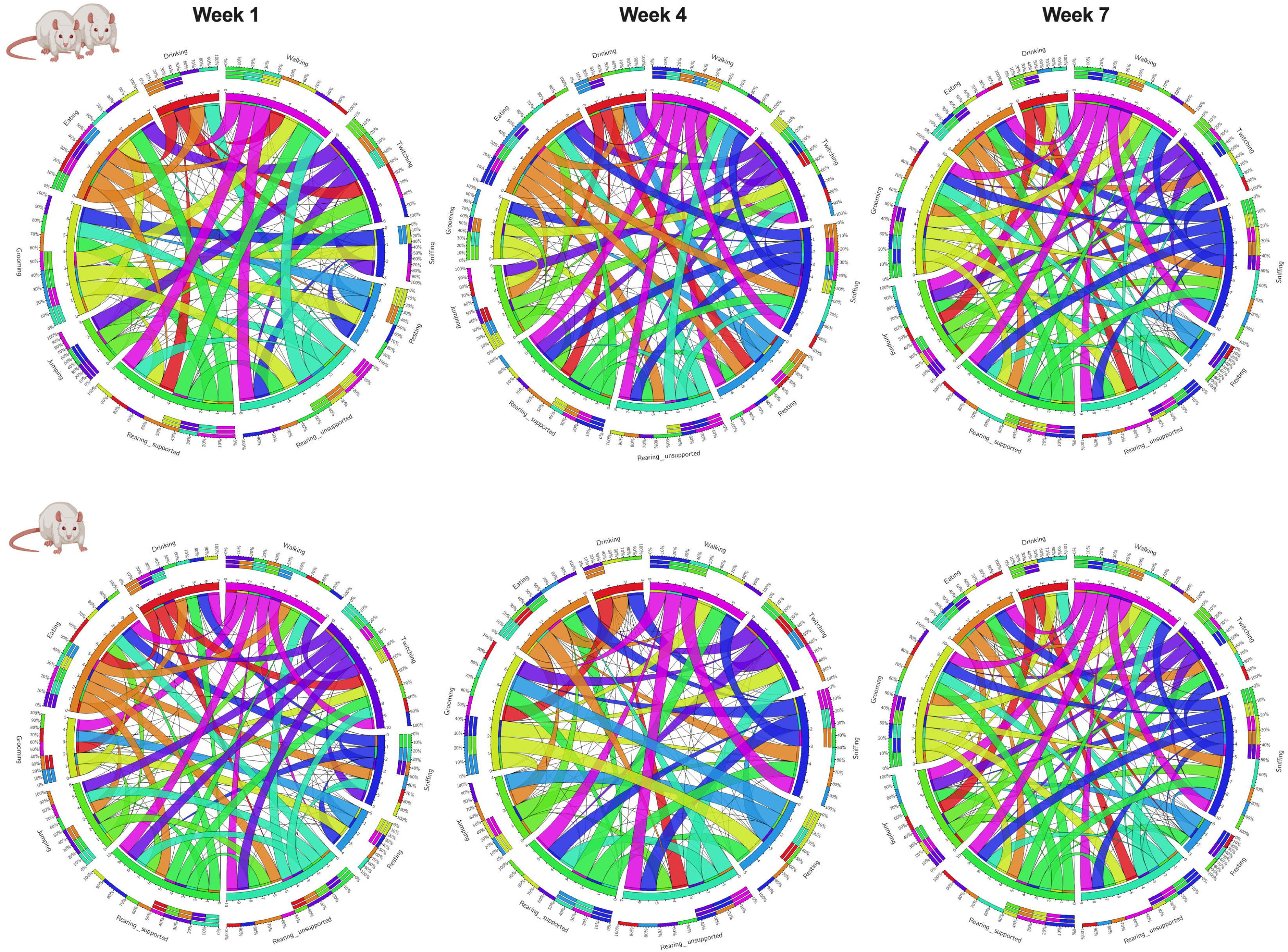
Organism-level Ethological Associations. This study evaluated a naturalistic behaviorome of Paired and Isolated animals and then connected these behaviors with emotionality, coping, and hormone levels. The connectogram 1. illustrates the feasibility of associating organism-level behaviors monitored through time, and 2. Social isolation stress changes the way behaviors correlate without one another. A thinker ribbon designates a stronger association.

## Supporting information

Supplemental Resources

## Contributions

**TS:** planned, performed, analyzed, interpreted experiments, prepared illustrations, wrote manuscript; **JS:** conducted experiments, performed quality control, statistical analyses, and analyzed behavioral data **AW and POA:** conducted experiments; **GW, AR, and JH:** performed quality control, statistical analyses, and analyzed behavioral data; **JL and FS:** carried out PhenoTyper experiments; **JDF:** planned experimental design and data interpretation, analyzed and interpreted data, wrote manuscript.

