## Supplemental Resources for "Shifts in Naturalistic Behaviors Induced by Early Social Isolation Stress are Associated with Adult Binge-like Eating in Female Rats"

**Supplemental Figures**

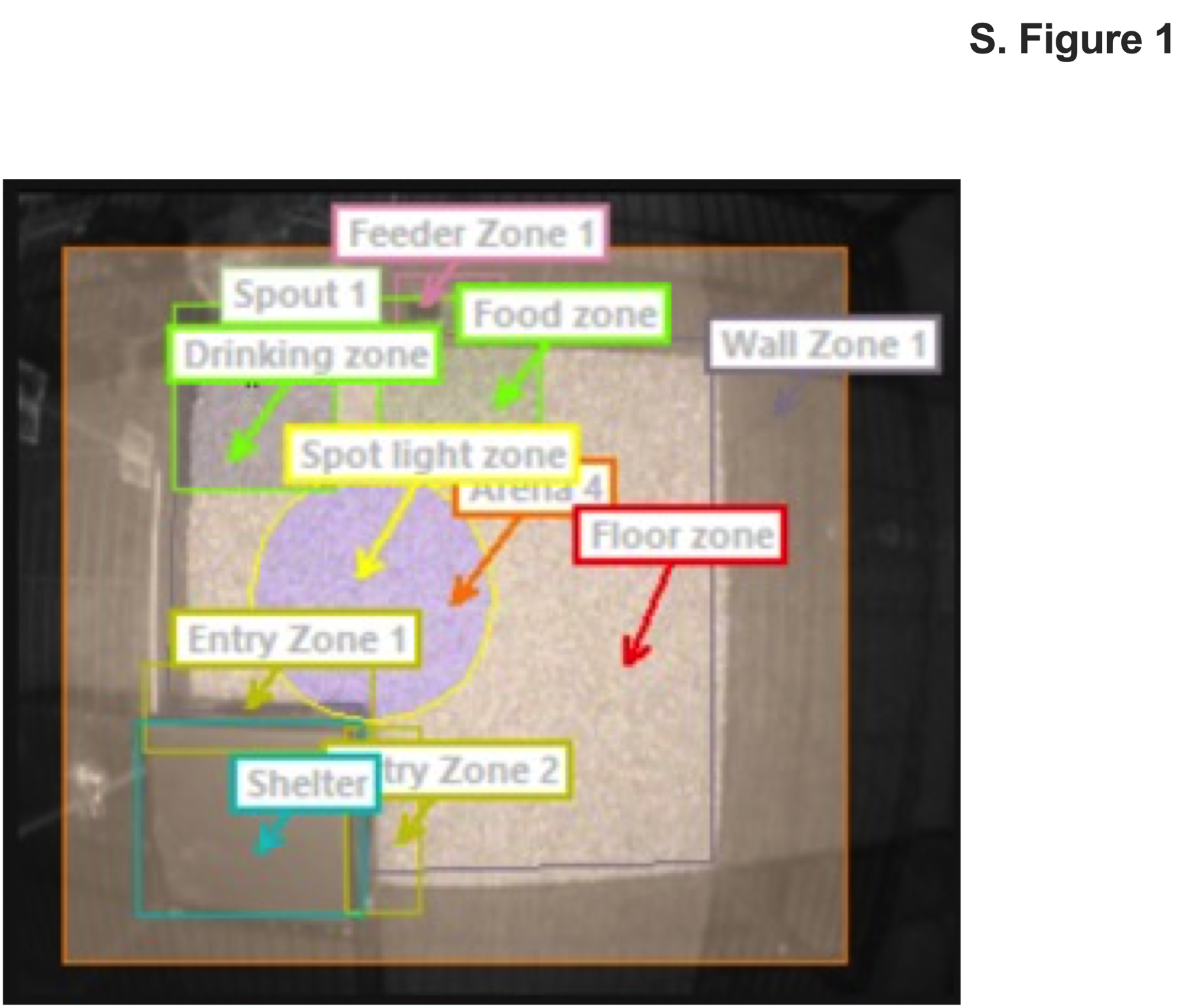
**S. Fig. 1.** *Aversive spotlight location and zones within the observational home cage*.

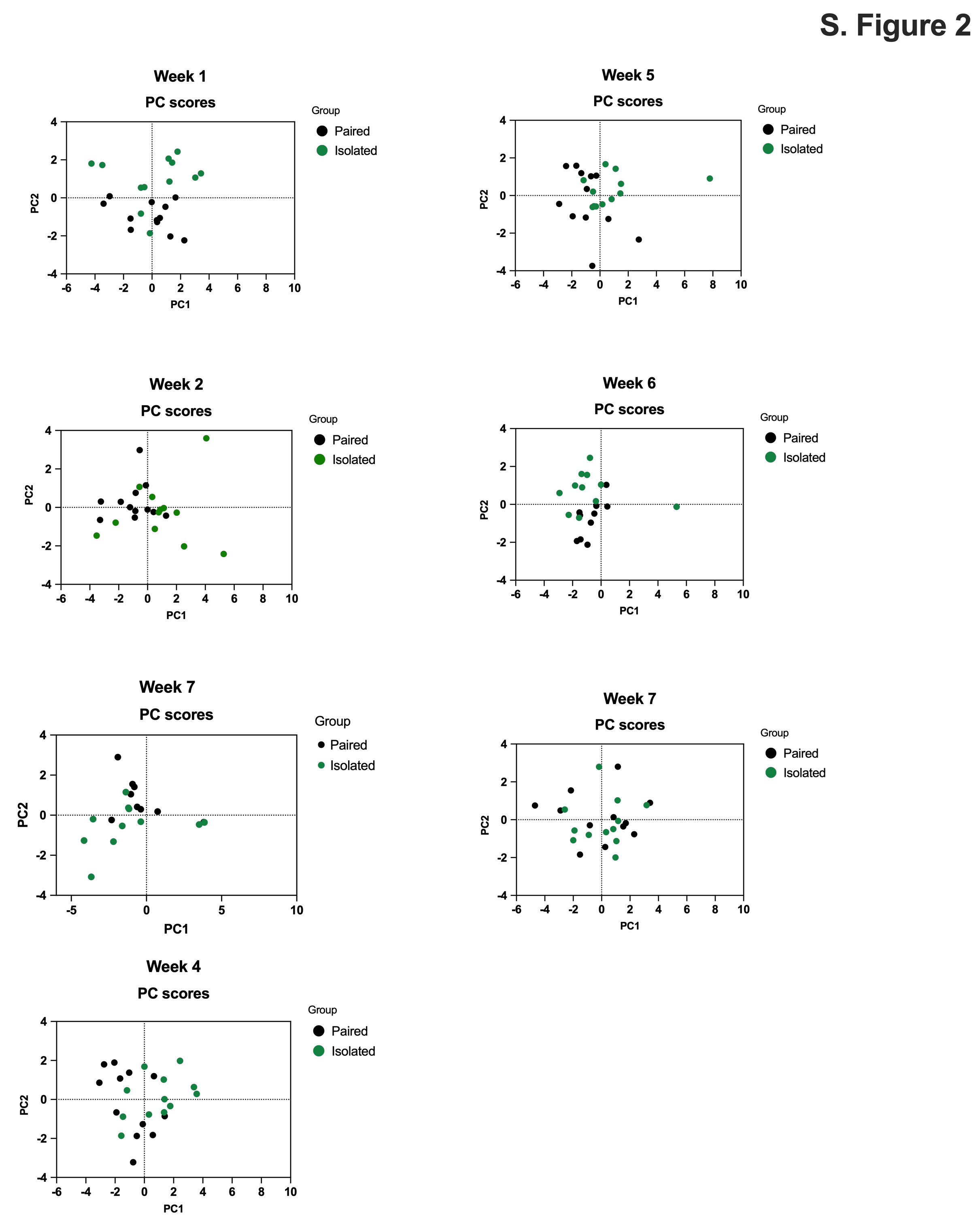
**S. Fig. 2.** *PCA of 10 naturalistic behaviors distinguishes Isolated and Paired animals.* The Paired animals in black, Isolated in green for all seven weeks. All the behaviors were standardized. Principal components were selected based off greater than 75% total explained variation.

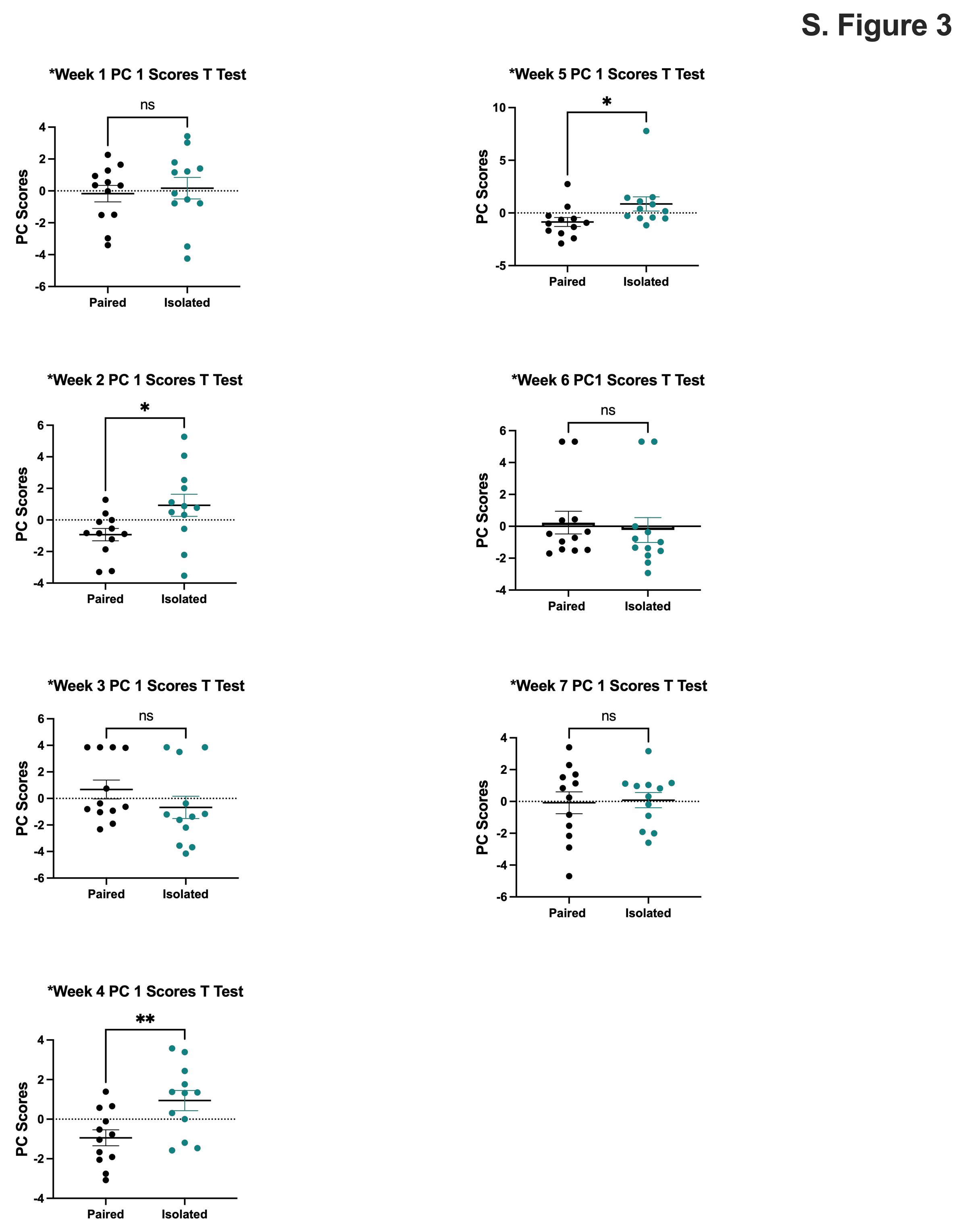
**S. Fig. 3.** *PC1 scores are altered in Isolated and Paired animals.* The Paired animals in black, Isolated in green for all seven weeks. Paired and Isolated, n=12 per group. Data presented with SEM. * p < 0.05, ** p < 0.01, *** p < 0.001, **** p <0.0001; Student t- tests.

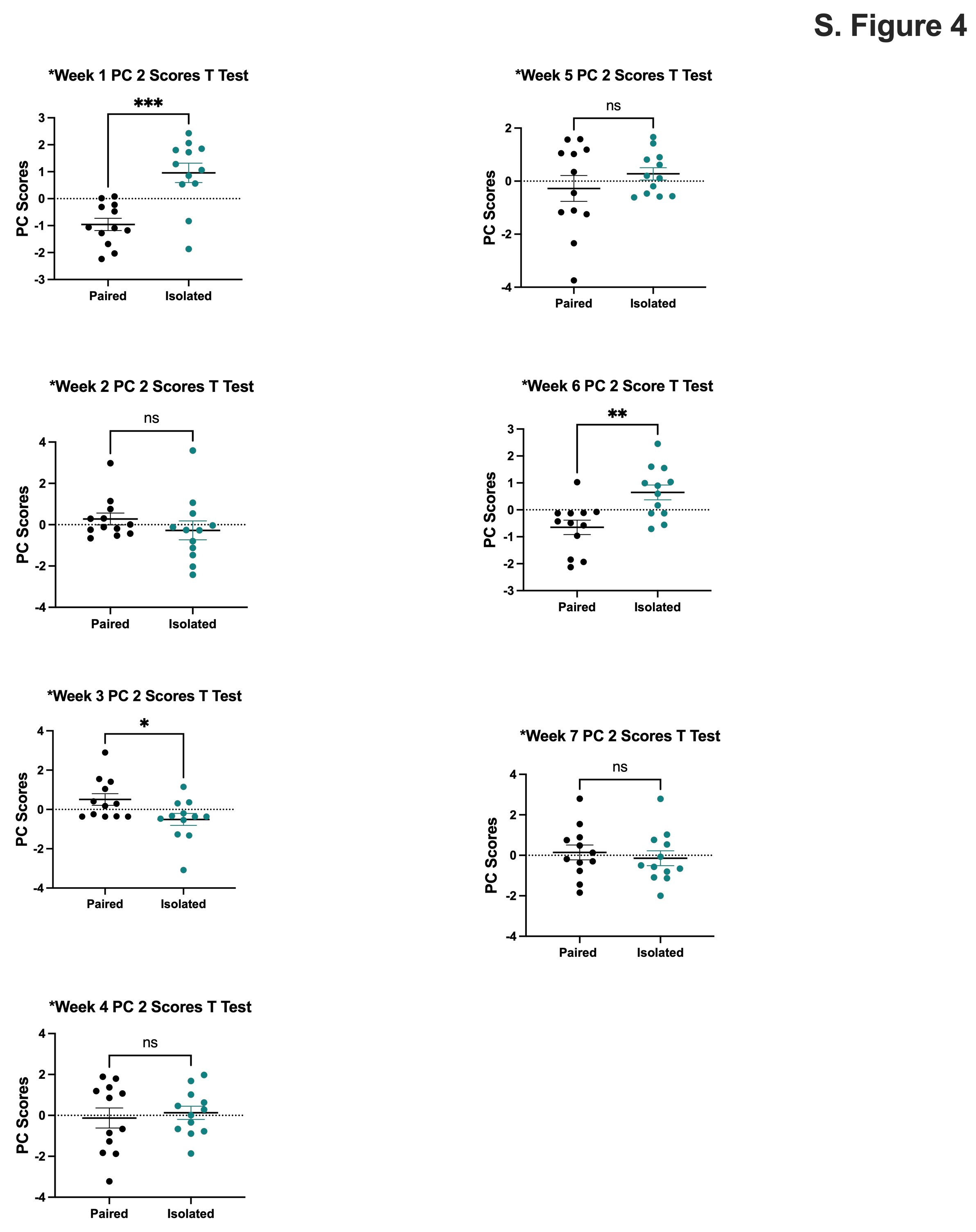
**S. Fig. 4.** *PC2 scores are altered in Isolated and Paired animals.* The Paired animals in black, Isolated in green for all seven weeks. Paired and Isolated, n=12 per group. Data presented with SEM. * p < 0.05, ** p < 0.01, *** p < 0.001, **** p <0.0001; Student t- tests.

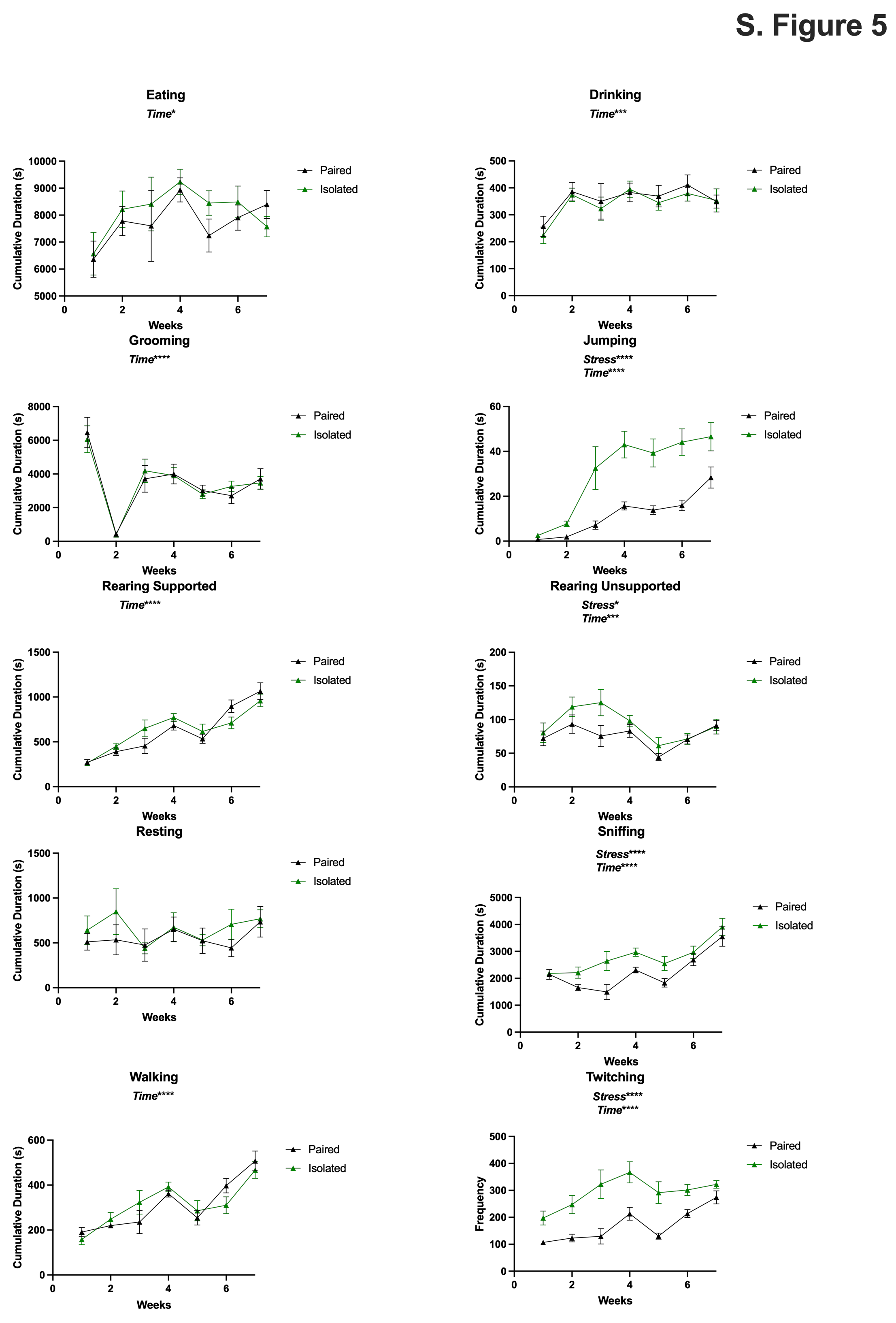
**S. Fig. 5.** *Weekly trajectories for each naturalistic behavior for all seven weeks.* Paired are in black, Isolation in green. Data presented with SEM. Paired and Isolated, n=12 per group. * p < 0.05, ** p < 0.01, *** p < 0.001, **** p <0.0001; Two-Way ANOVA.

**S. Fig. 6.** *Shifted behaviors capture the phenotypic profile of SI animals.* **(A)** Shifted behaviors include Rearing Unsupported, Jumping, Sniffing, and Twitching. **(B)** Nonshifted included the remaining six behaviors. Paired and Isolated, n=12. Data presented with SEM. * p < 0.05, ** p < 0.01, *** p < 0.001, **** p <0.0001; Student t- tests.

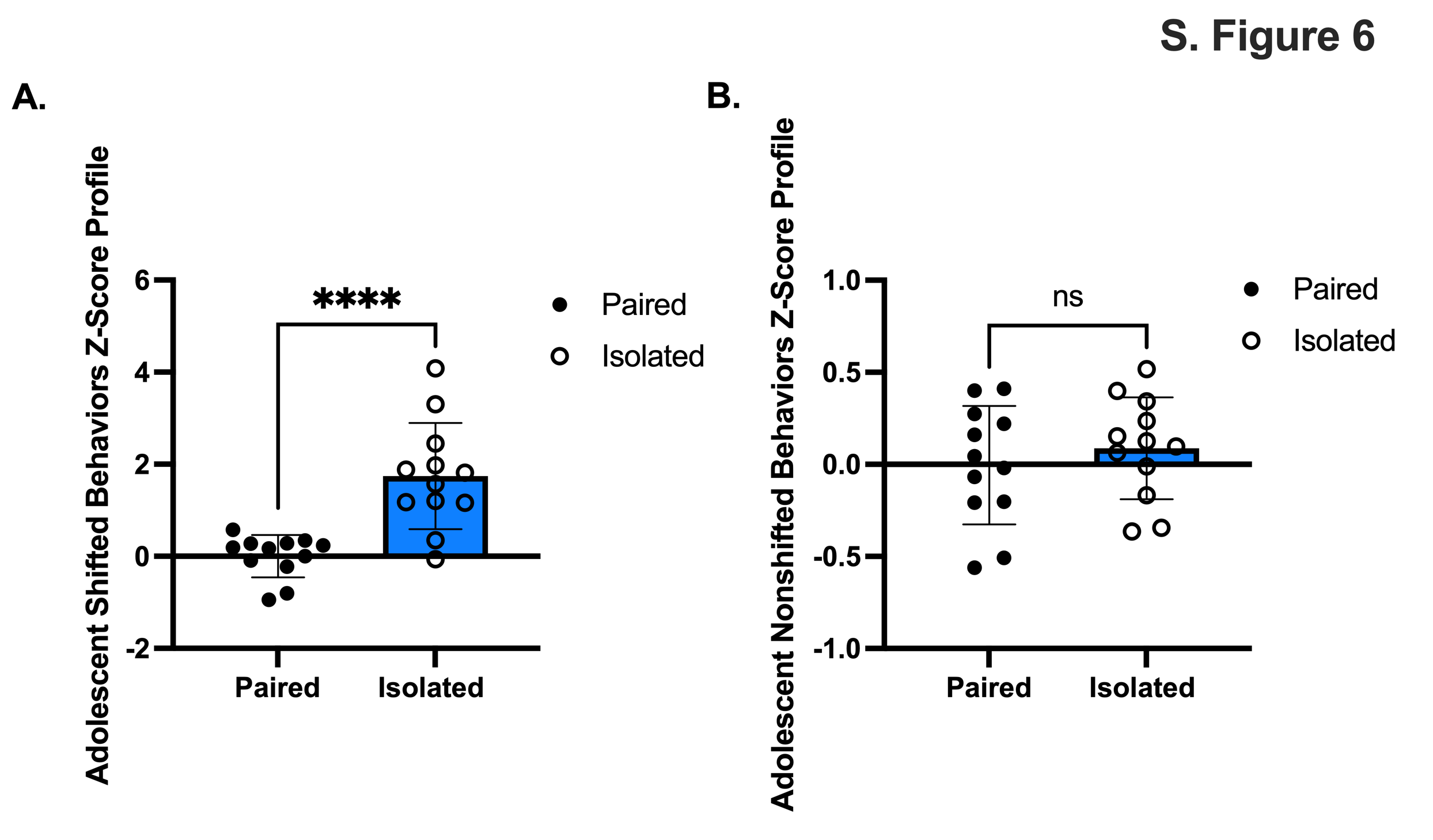

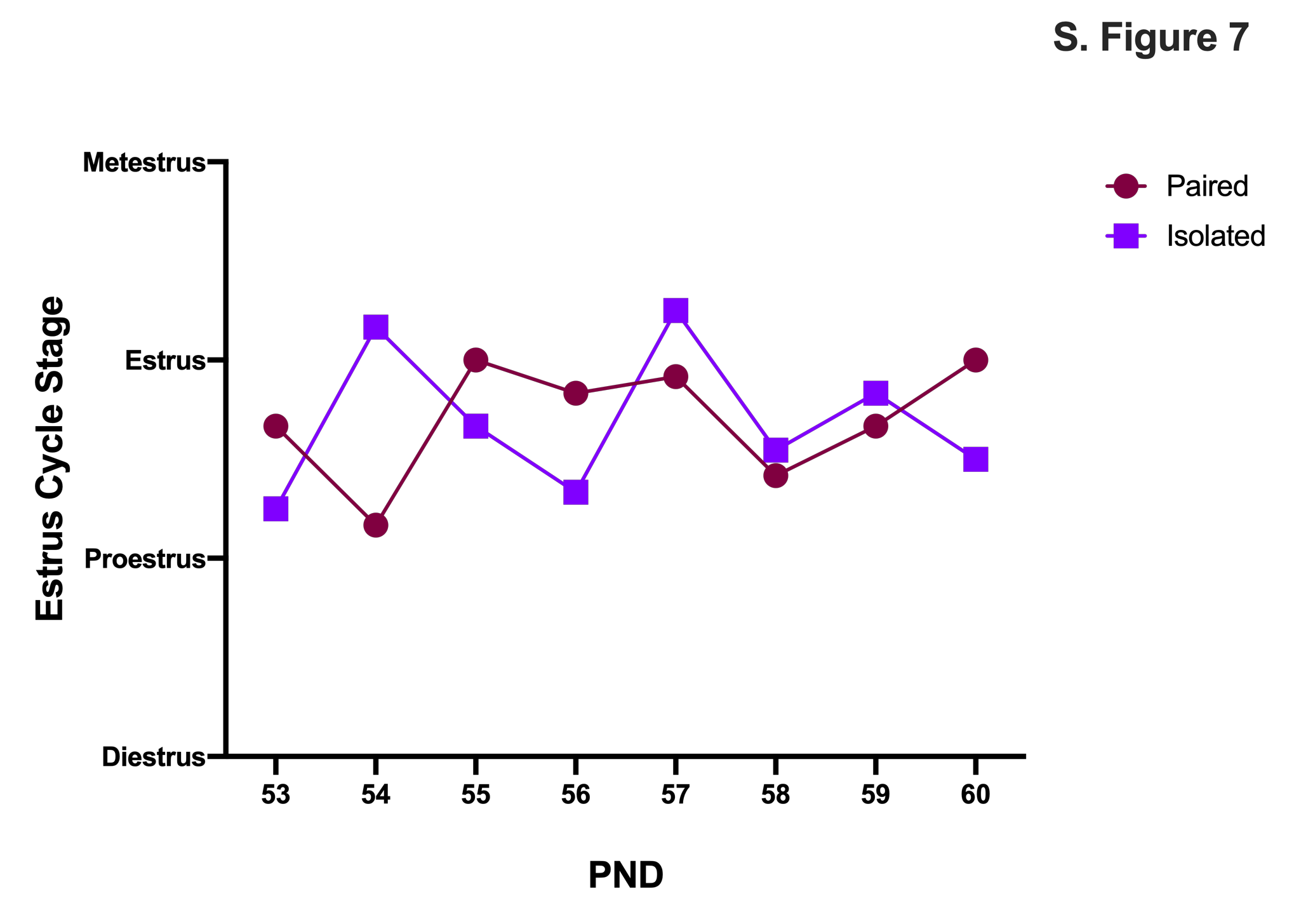
**S. Fig. 7.** *Estrus Cyclicity does not differ between Paired and Isolated groups during adolescence.*

**S. Fig. 8.** *An aversive spotlight challenge.* The dark grey rectangle depicts the start and stop of the spotlight hour. Each behavior is shown for all 7 weeks. Paired are in black, Isolation in green. Data presented with SEM. Paired and Isolated, n=12 per group. * p < 0.05, ** p < 0.01, *** p < 0.001, **** p <0.0001; Two-Way ANOVA.

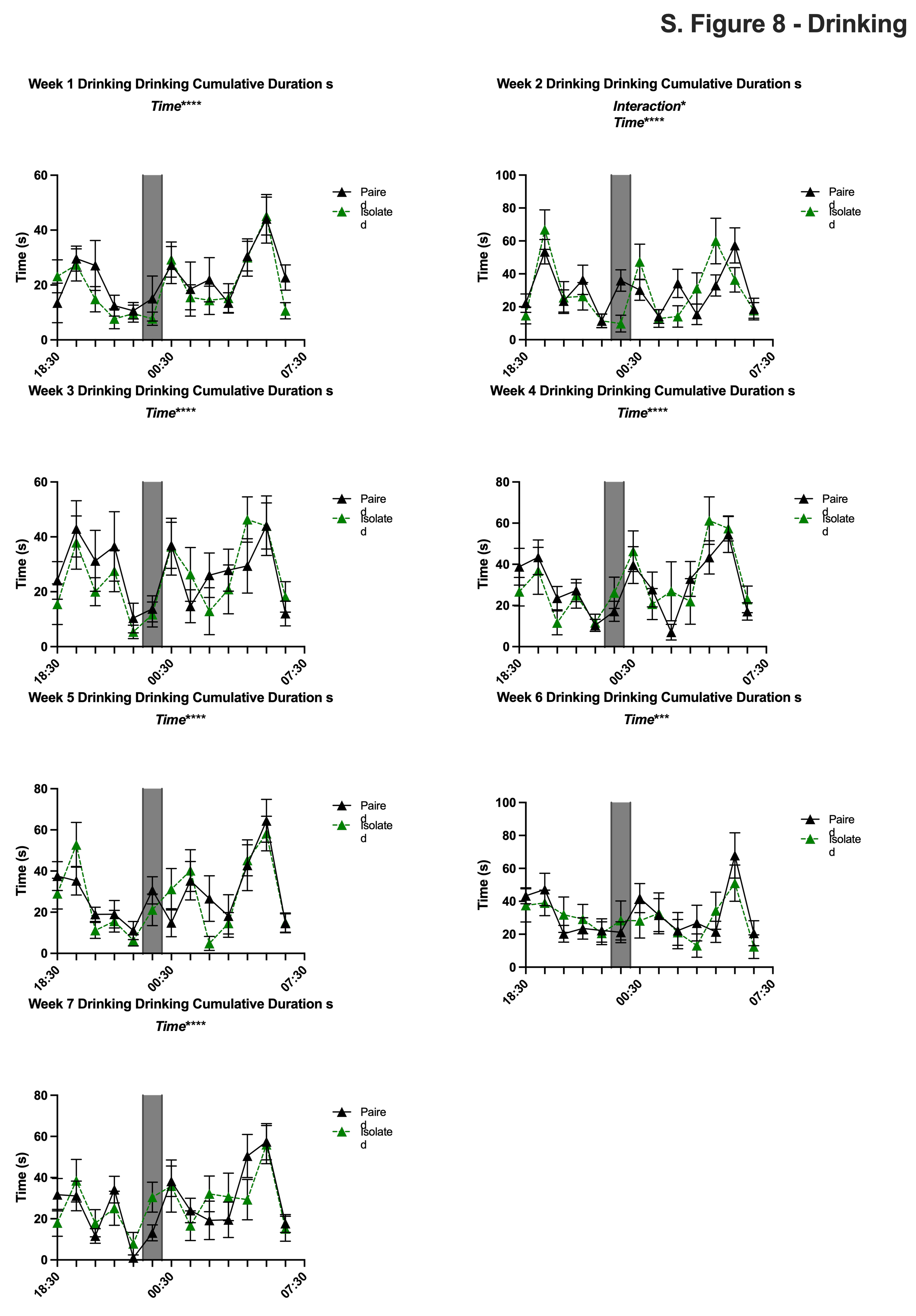

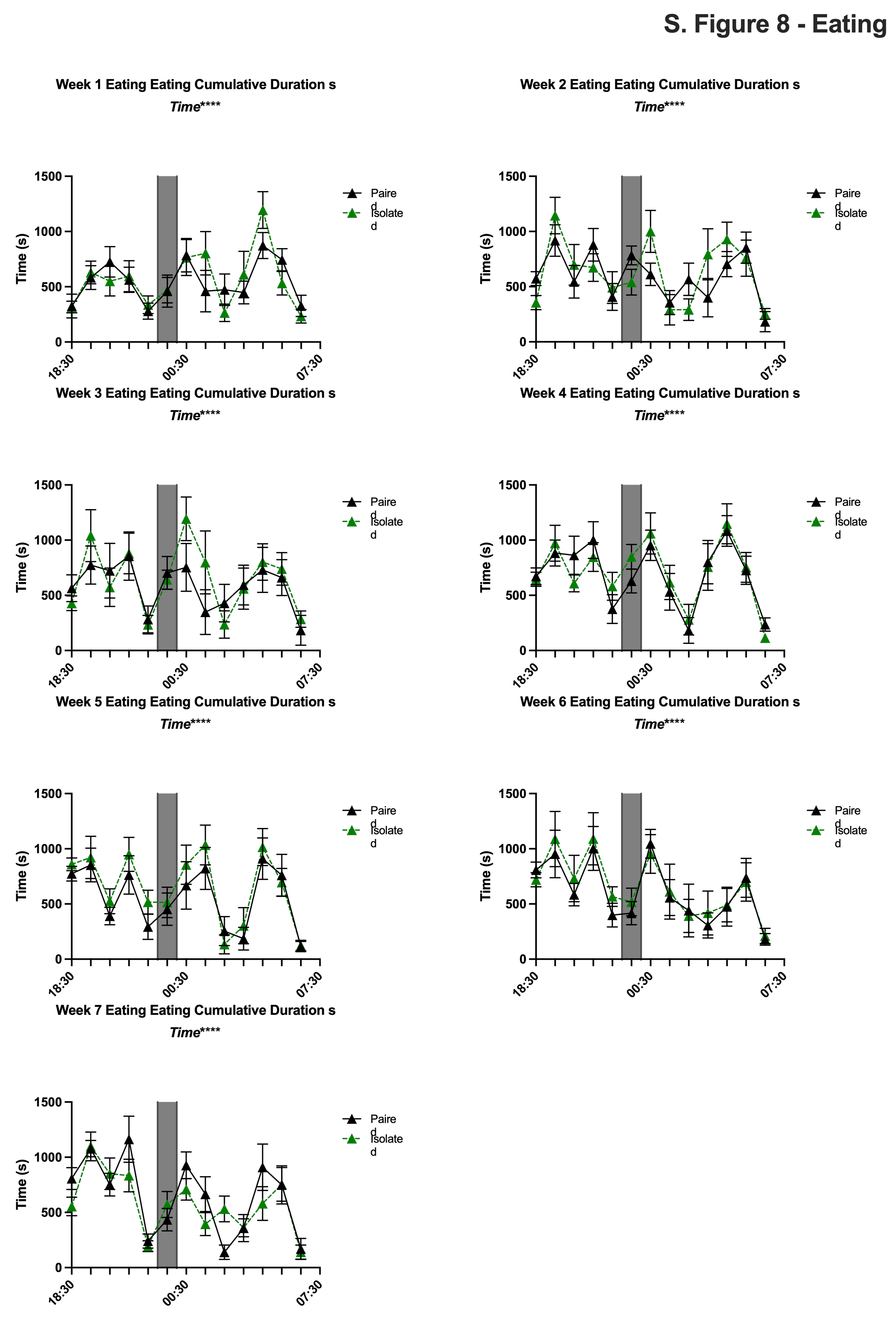

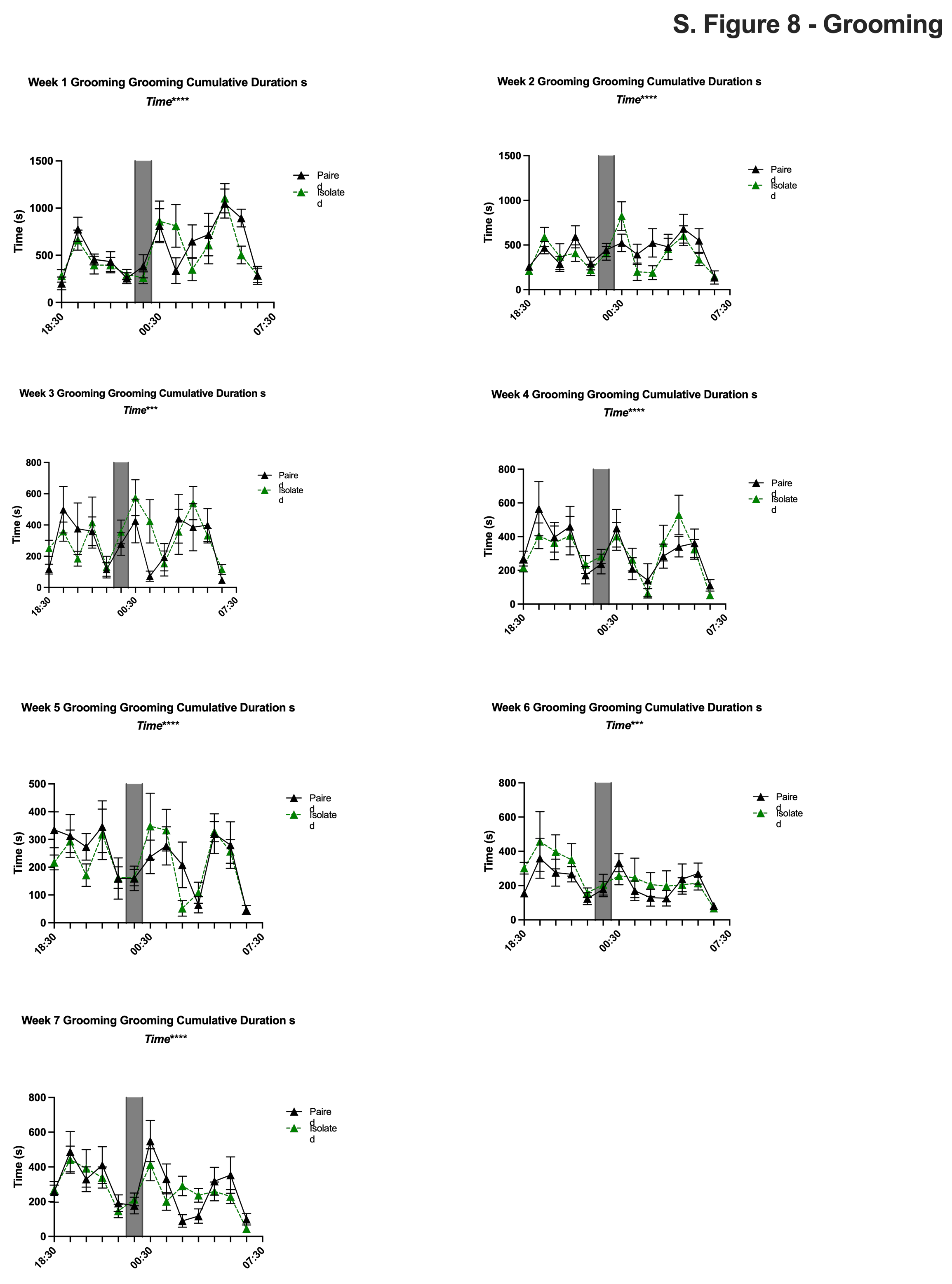

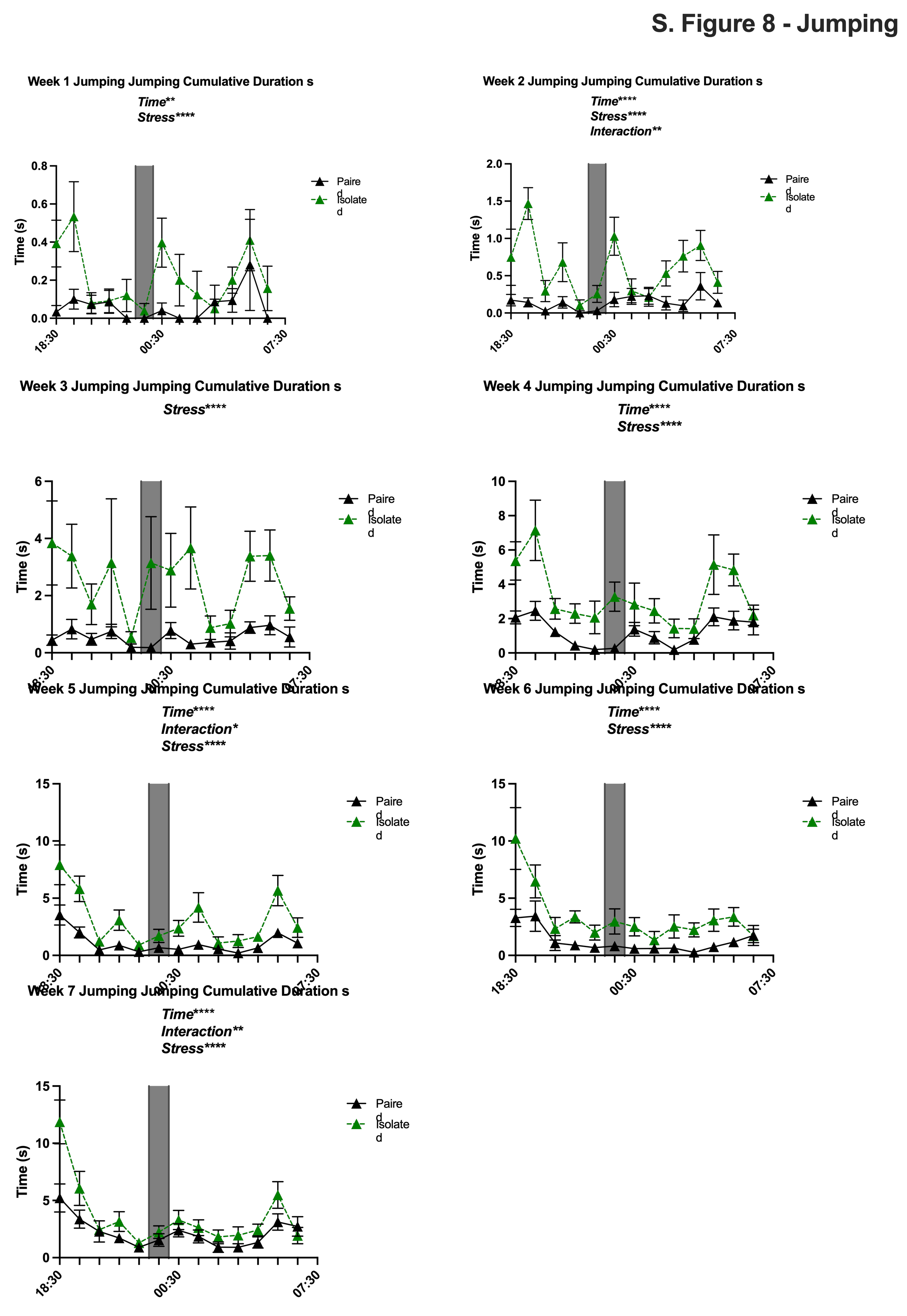

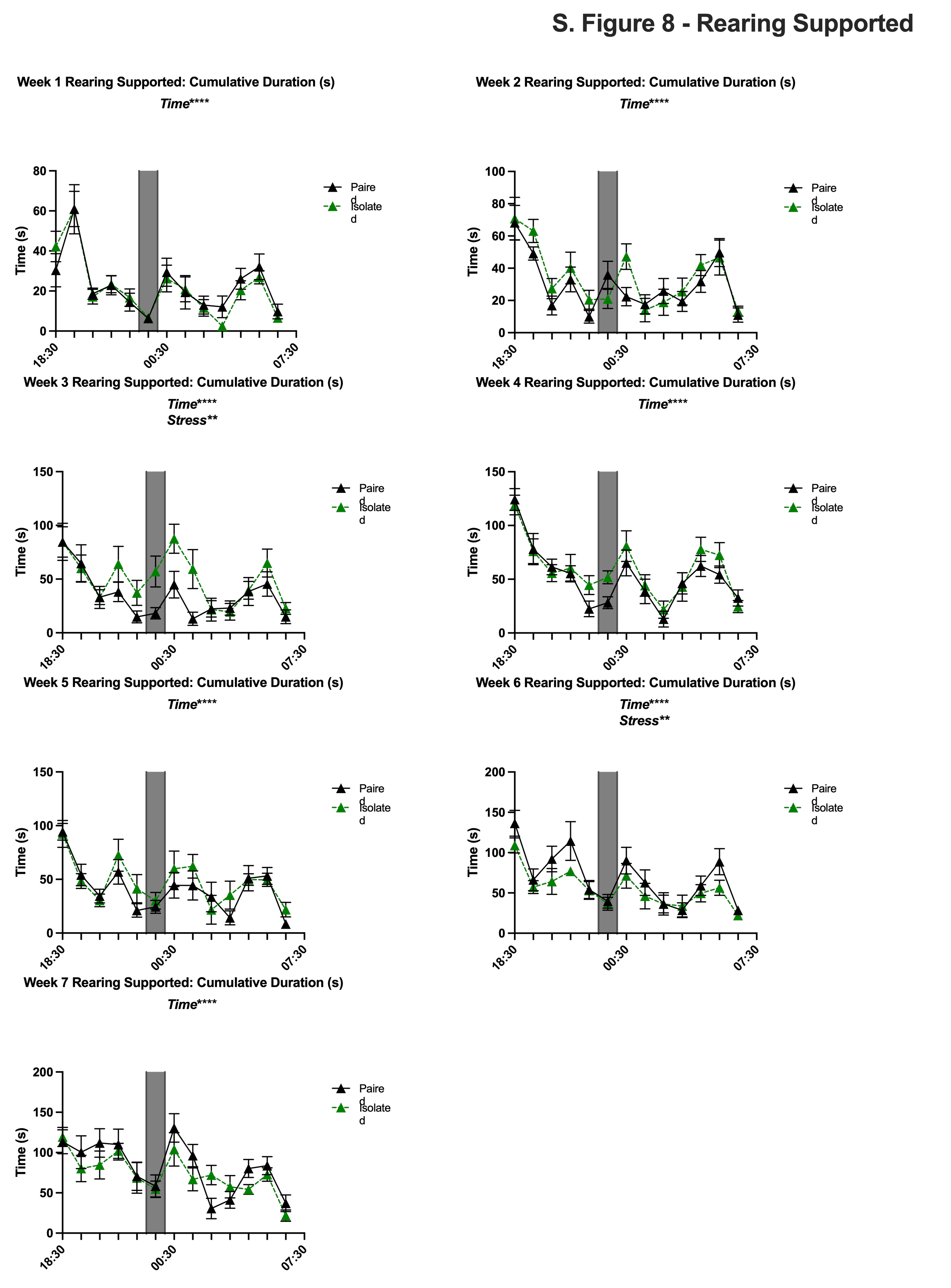

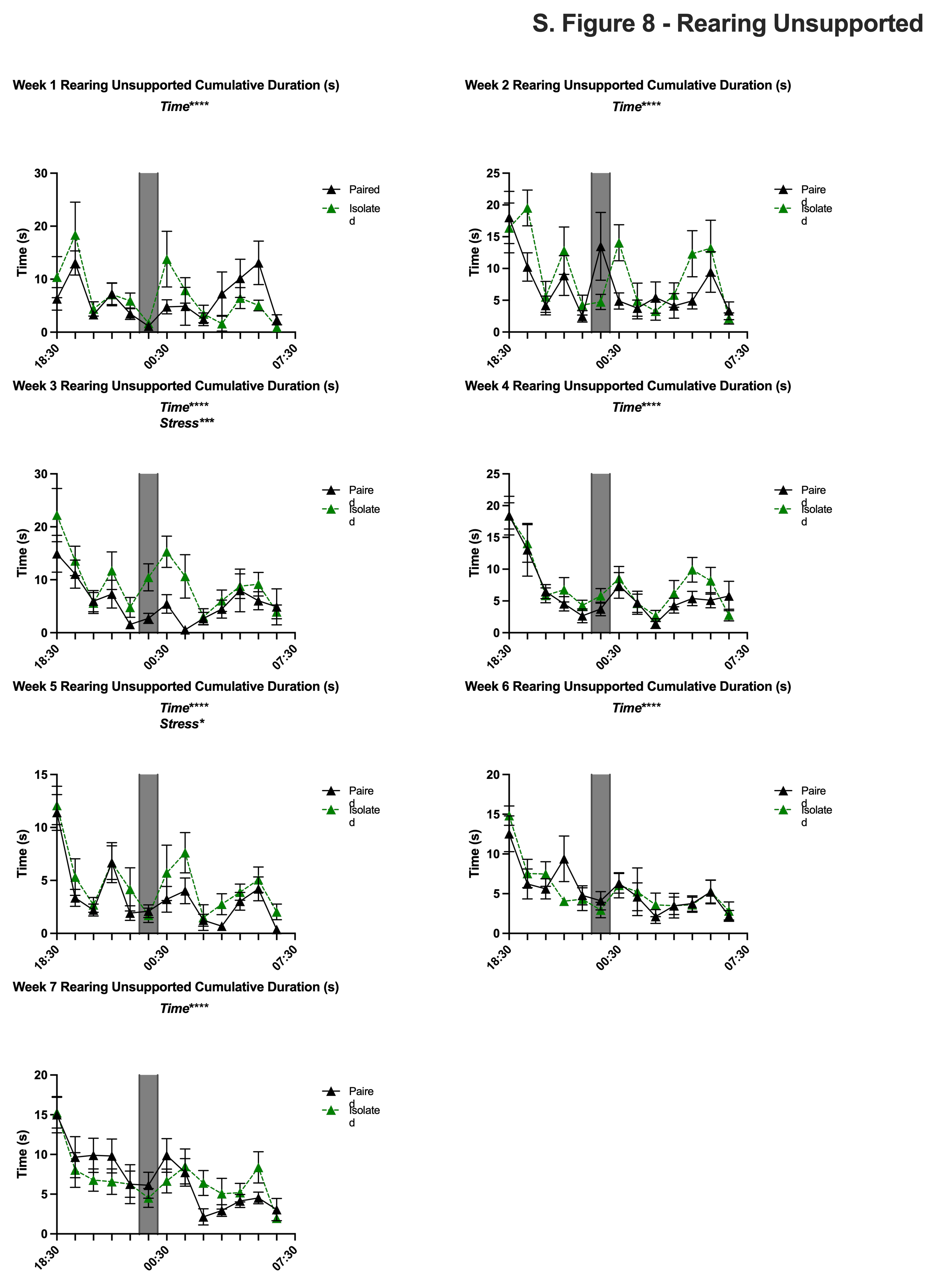

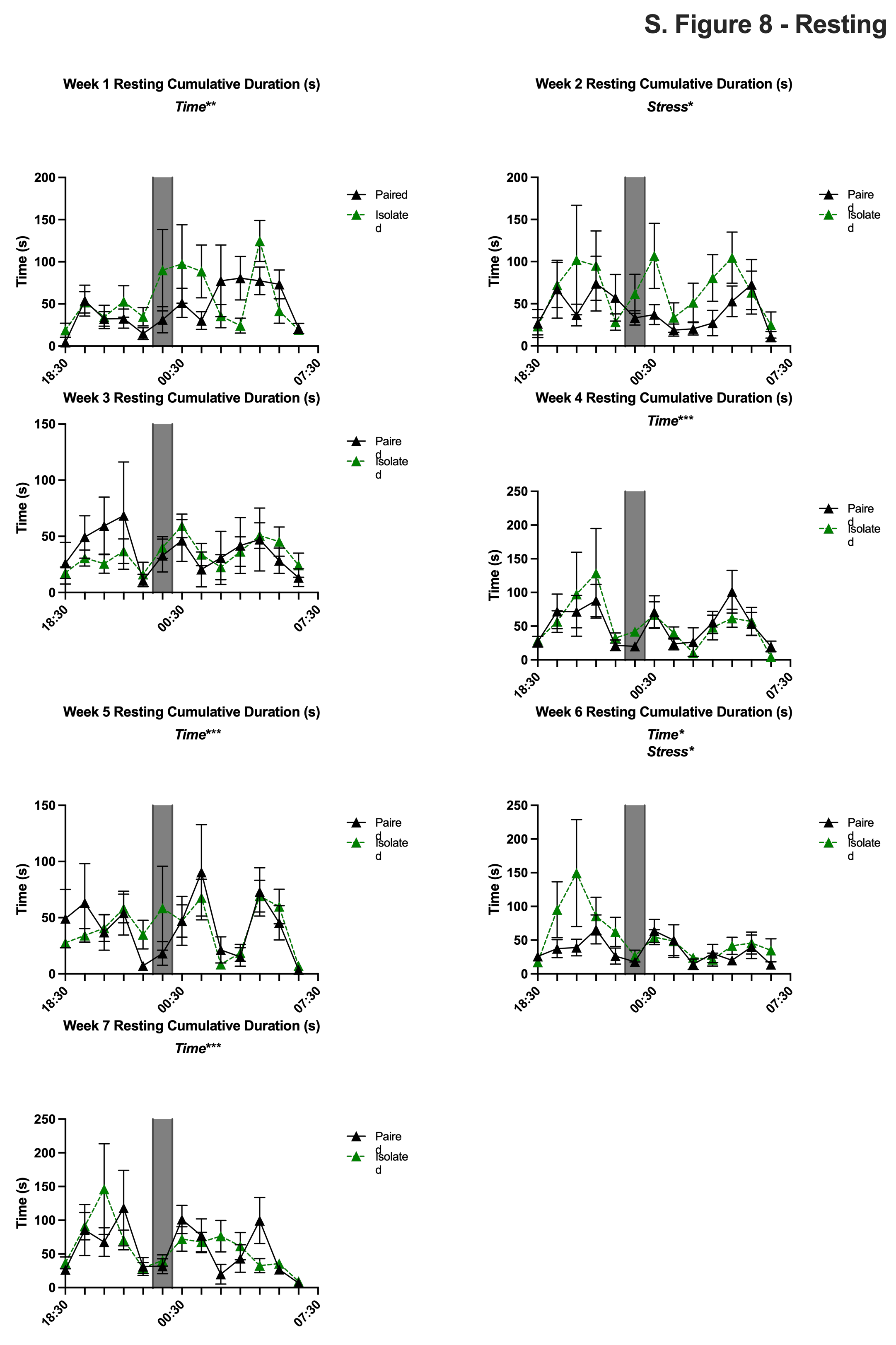

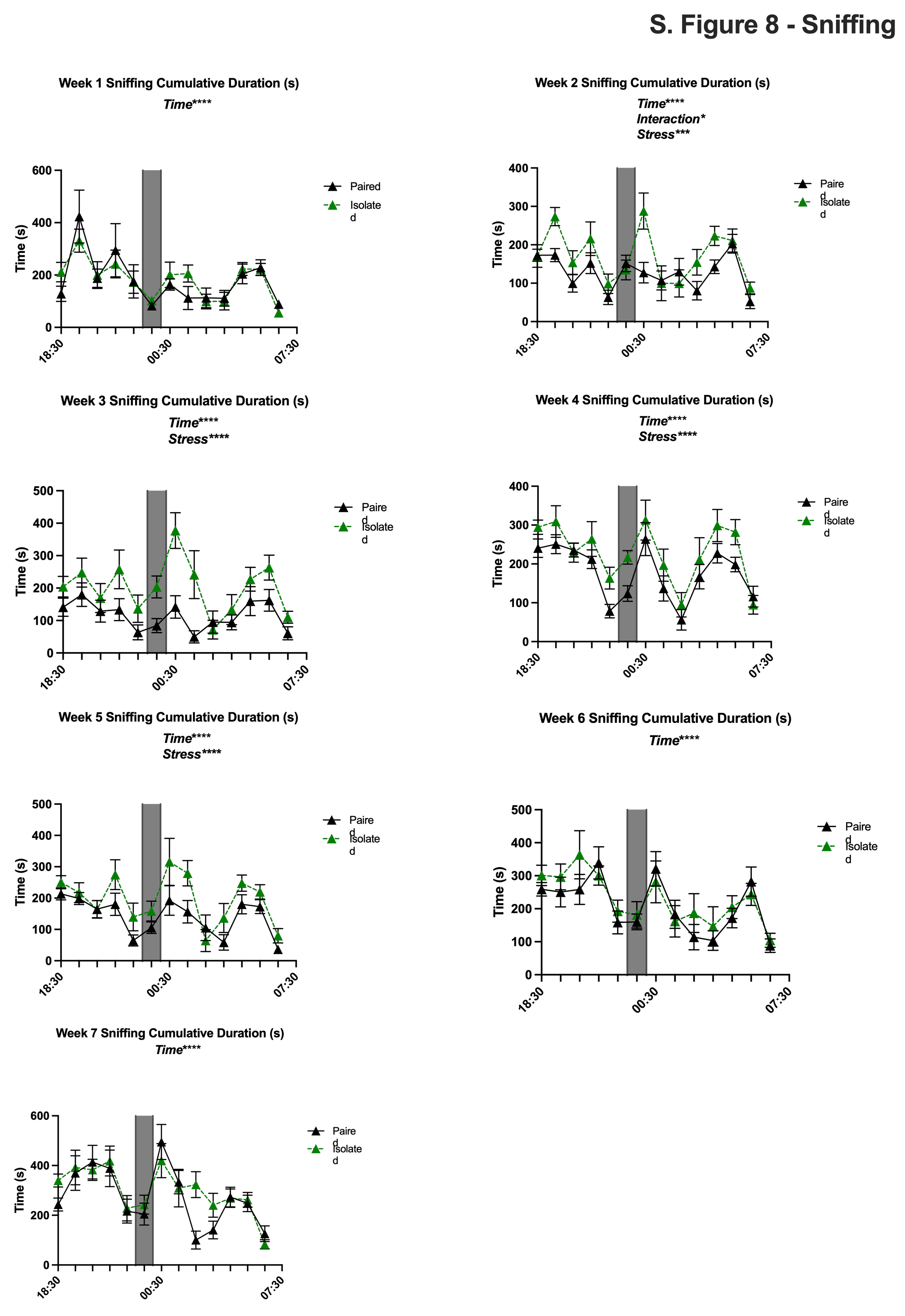

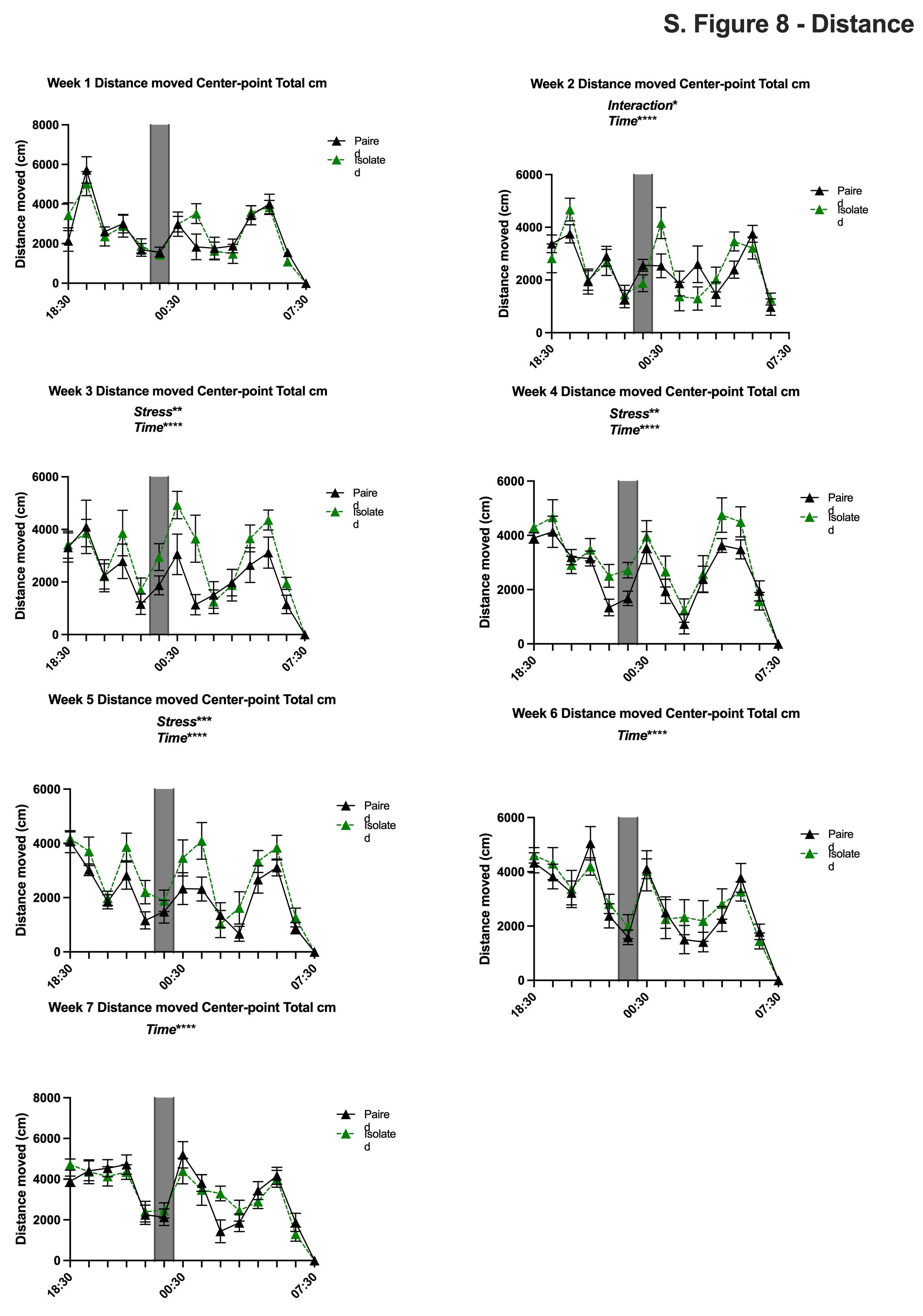

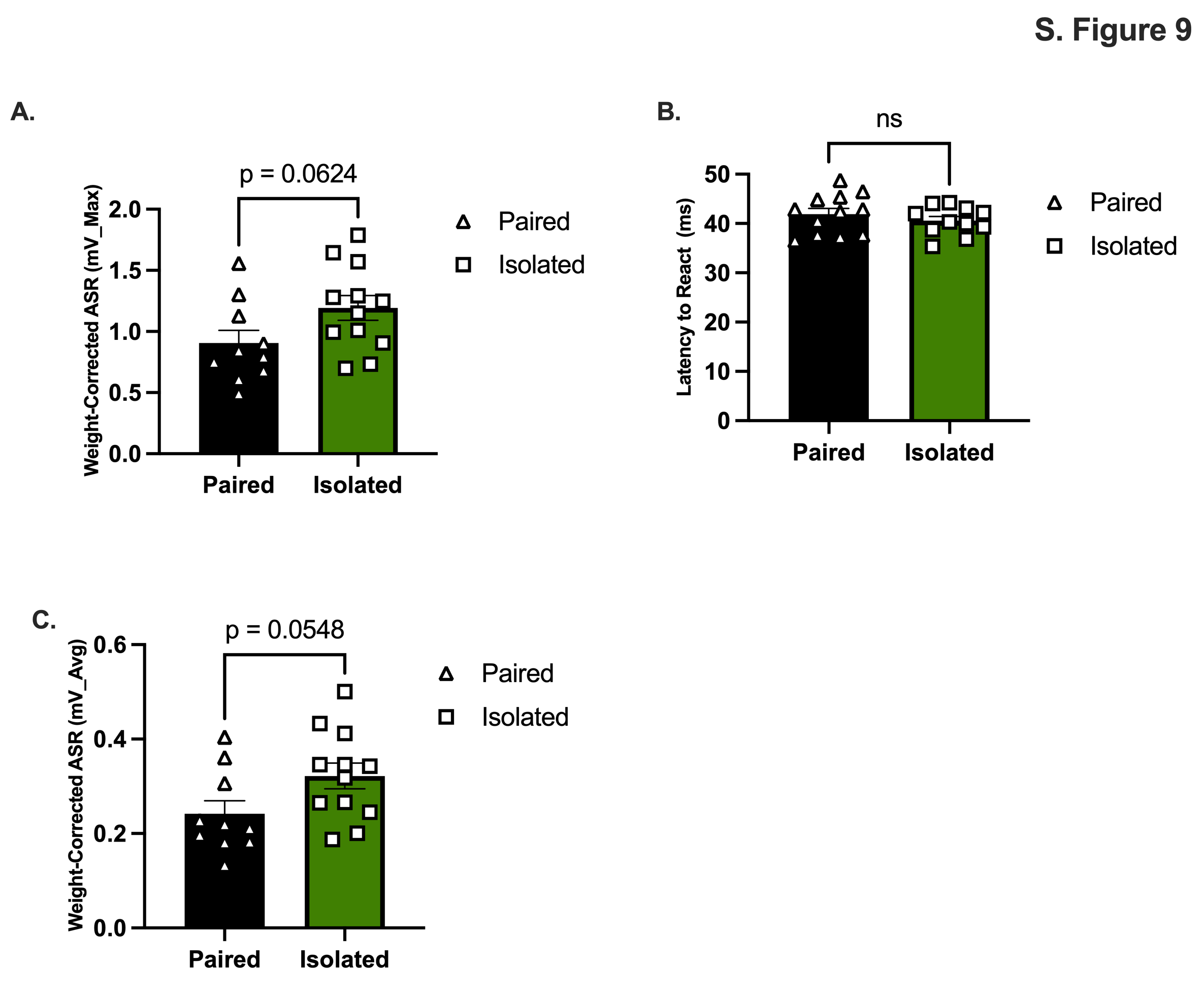
**S. Fig. 9.** *Acoustic startle Reactivity. Isolated animals hyper react compared to Paired counterparts.* **(A)** Weight-corrected maximum amplitude. **(B)** Latency to react in milliseconds. **(C)** Weight-corrected average amplitude. Data presented with SEM. * p < 0.05, ** p < 0.01, *** p < 0.001, **** p <0.0001; Student t- tests for parametric and Mann-Whitney for nonparametric.

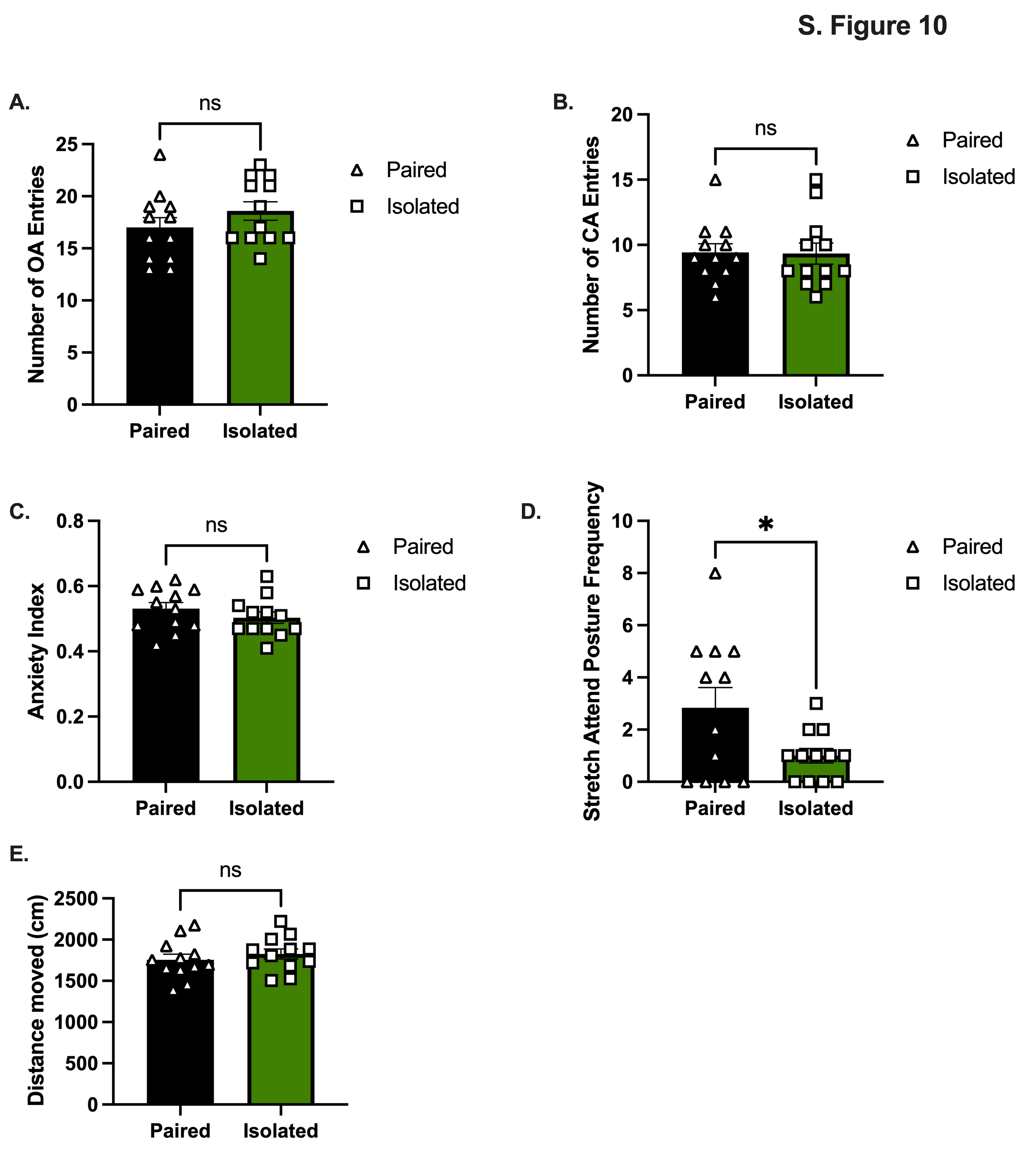
**S. Fig. 10.** *Elevated Plus Maze.* **(A, B, C, and E)** No differences in EPM. **(D)** Stretch attend posture is significantly decreased in Isolated animals. OA = open arms; CA = closed arms. Data presented with SEM. * p < 0.05, ** p < 0.01, *** p < 0.001, **** p <0.0001; Student t- tests for parametric and Mann-Whitney for nonparametric.

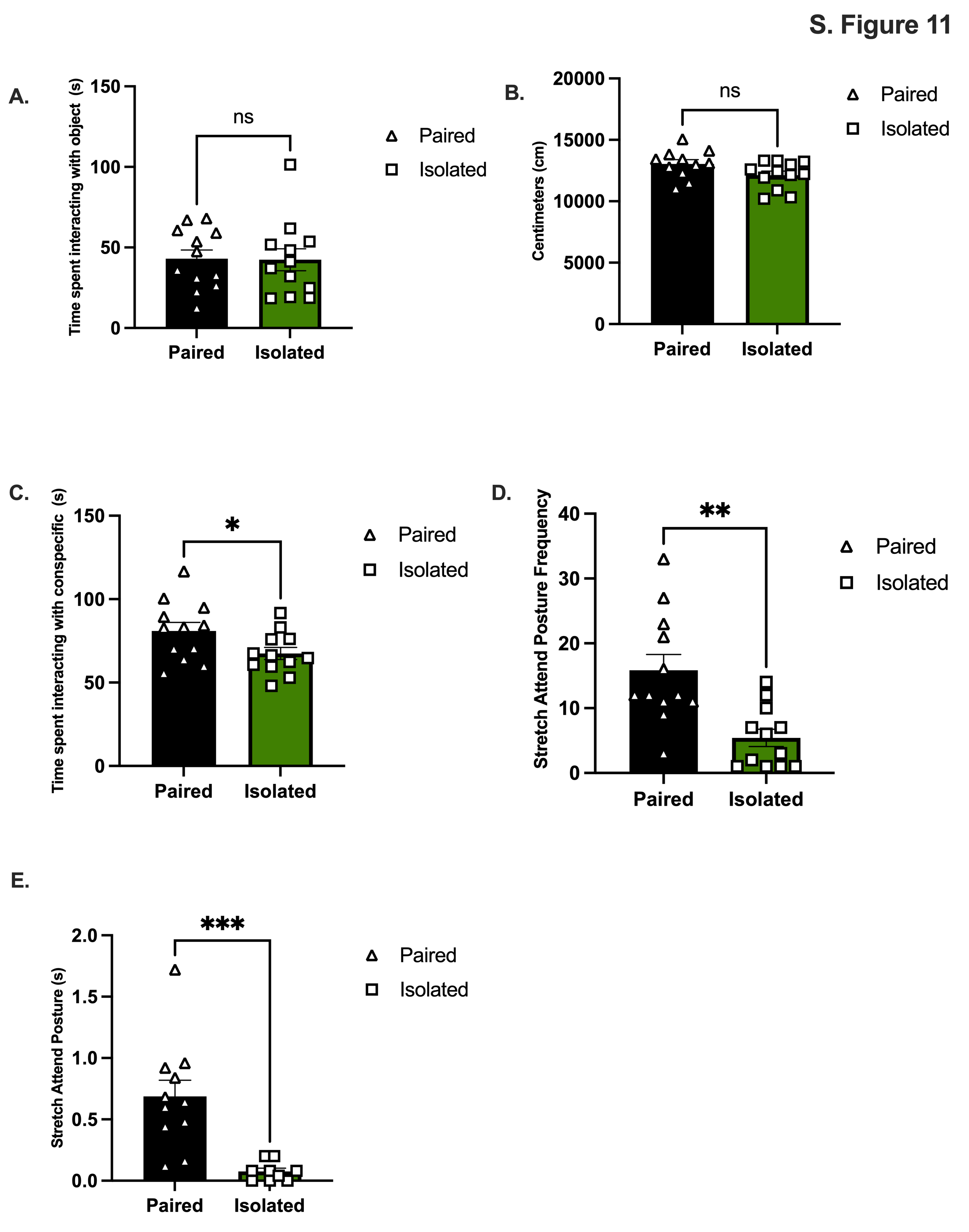
**S. Fig. 11.** *Social Y Maze.* *Isolated have decreased sociability compared to Paired counterparts.* **(A)** Time spent with the object. **(B)** the distance traveled in centimeters. **(C)** Time spent interacting with the age and sex matched conspecific. **(D)** Number of times the rats stretch attended. **(E)** Stretch attend duration in seconds. Data presented with SEM. * p < 0.05, ** p < 0.01, *** p < 0.001, **** p <0.0001; Student t- tests for parametric and Mann-Whitney for nonparametric.

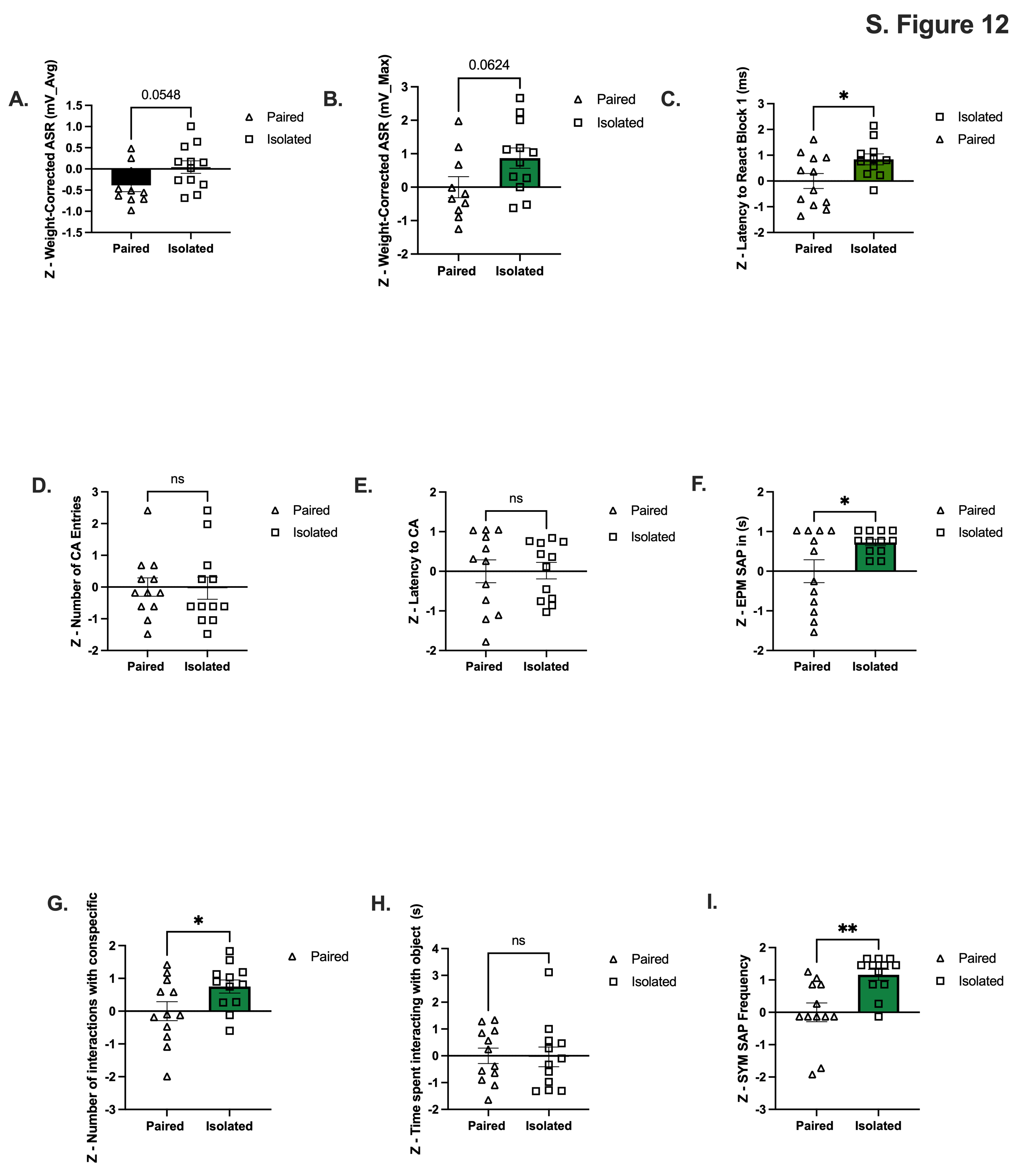
**S. Fig. 12.** *Metrics used to calculate emotionality.* Metrics with the highest contribution to variation based off PCA were chosen for emotionality scoring. Each metric was converted to a z-score prior to being averaged. See main figure 4. **(A-C)** ASR metrics include weight corrected average and maximum amplitude, and latency to react during the first block (10 trials). **(D-F)** EPM metrics include the number CA entries, latency to enter the CA, and duration of stretch attend postures. CA = closed arm. **(G-I)** SYM metrics include the number of interactions with the conspecific, time spent with object and stretch attend frequency. Data presented with SEM. * p < 0.05, ** p < 0.01, *** p < 0.001, **** p <0.0001; Student t- tests.

**S. Fig. 13.** *Generating a social isolation-induced binge-like eating model.* Lewis rats underwent chronic social isolation throughout adolescence prior to receiving WD in adulthood, n=15-16 per group. Data presented with SEM* p < 0.05, Two-Way ANOVA and Tukey HSD post hoc.

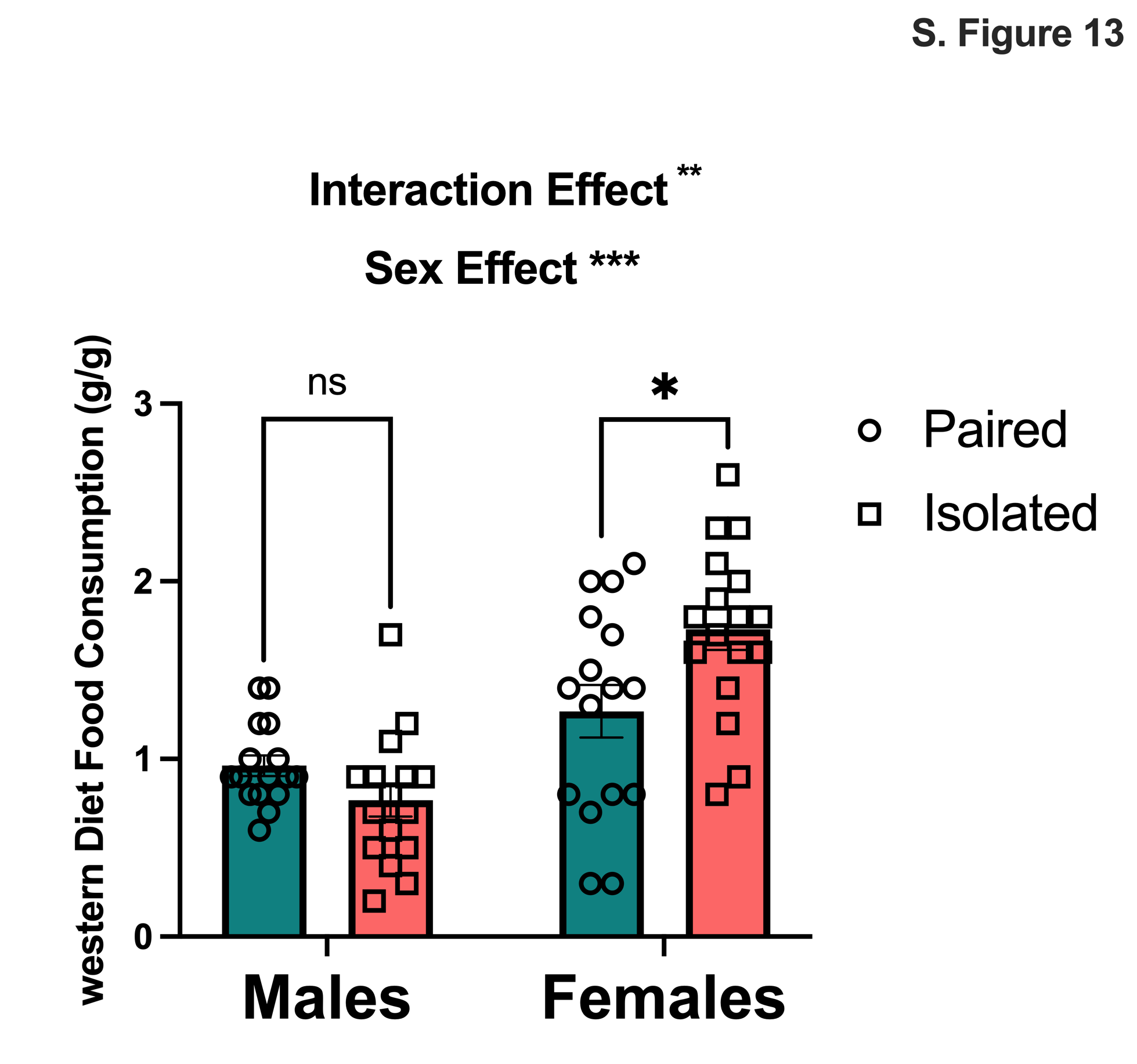

**S. Fig. 14.** *Estrus cycle hormones.* **(A-C)** There were no detectable differences in estrus cycle hormone levels in plasma, n=11-12. **(D)** PCA analysis with the estrus hormones further confirms overlap between groups. Data presented with SEM. Student t- tests.

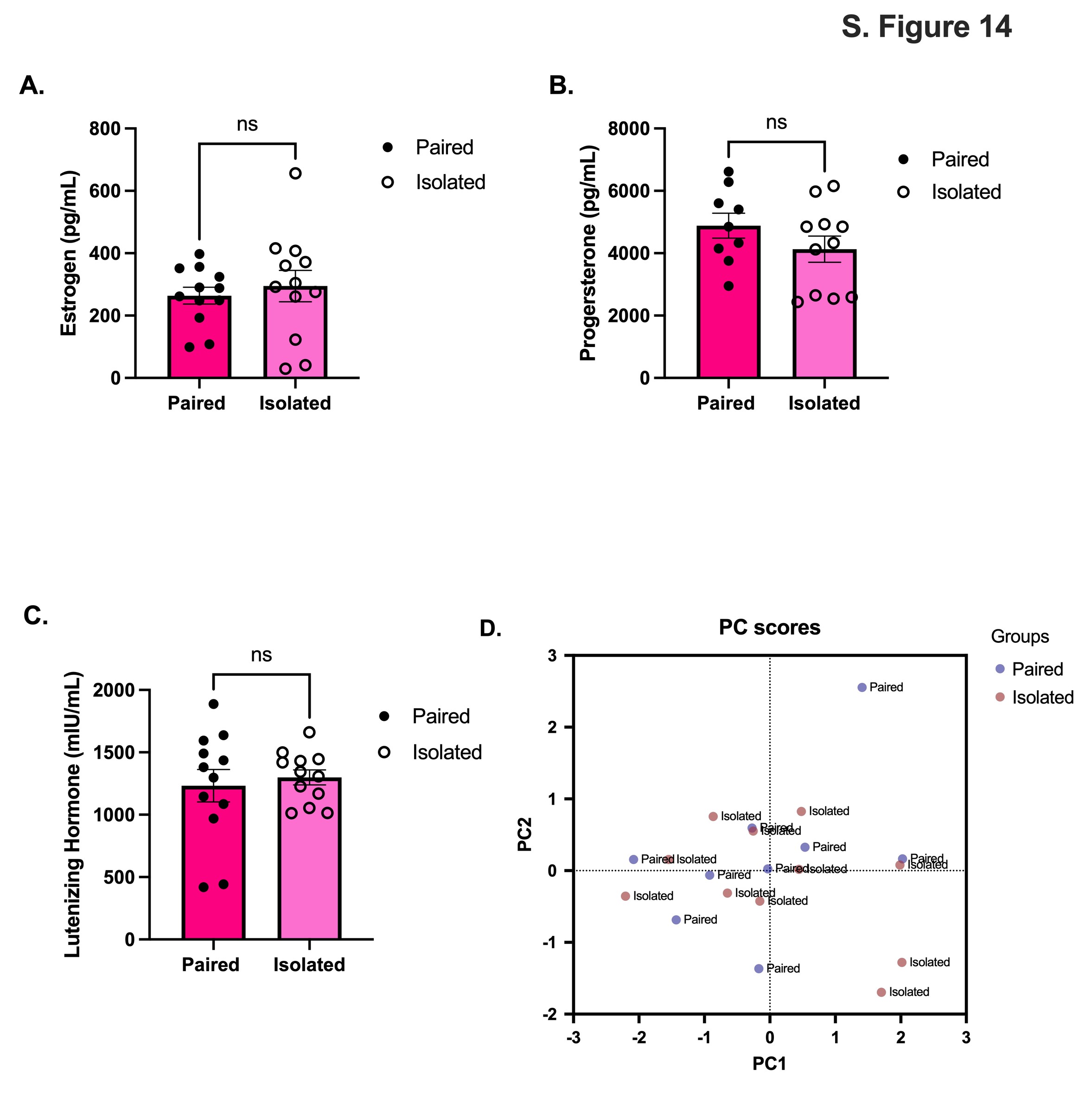

**S. Fig. 15.** *Naturalistic behaviors correlate with estrus cycle hormones.* **(A)** Estrogen and **(B)** Progesterone were correlated with Rearing Unsupported. **(C)** Progesterone was correlated with plasma levels of corticosterone. When separating Paired and Isolated groups, **(D)** only estrogen levels in the Paired animals remained correlated with Rearing Unsupported. **(E)** Isolated rat’s progesterone correlated with Rearing Unsupported. * p < 0.05, ** p < 0.01; Pearson Correlations conducted for each comparison. p and r values are presented.

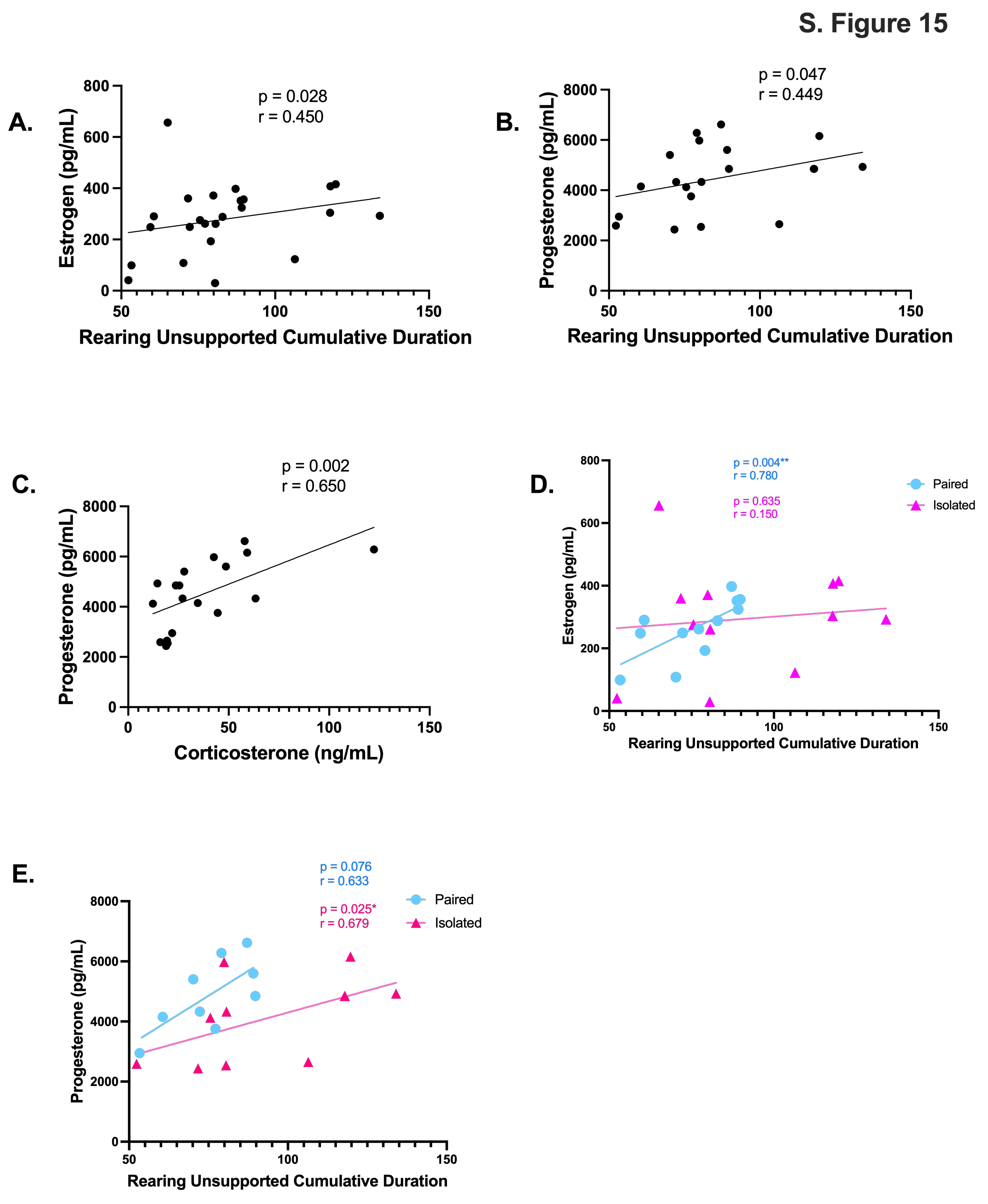

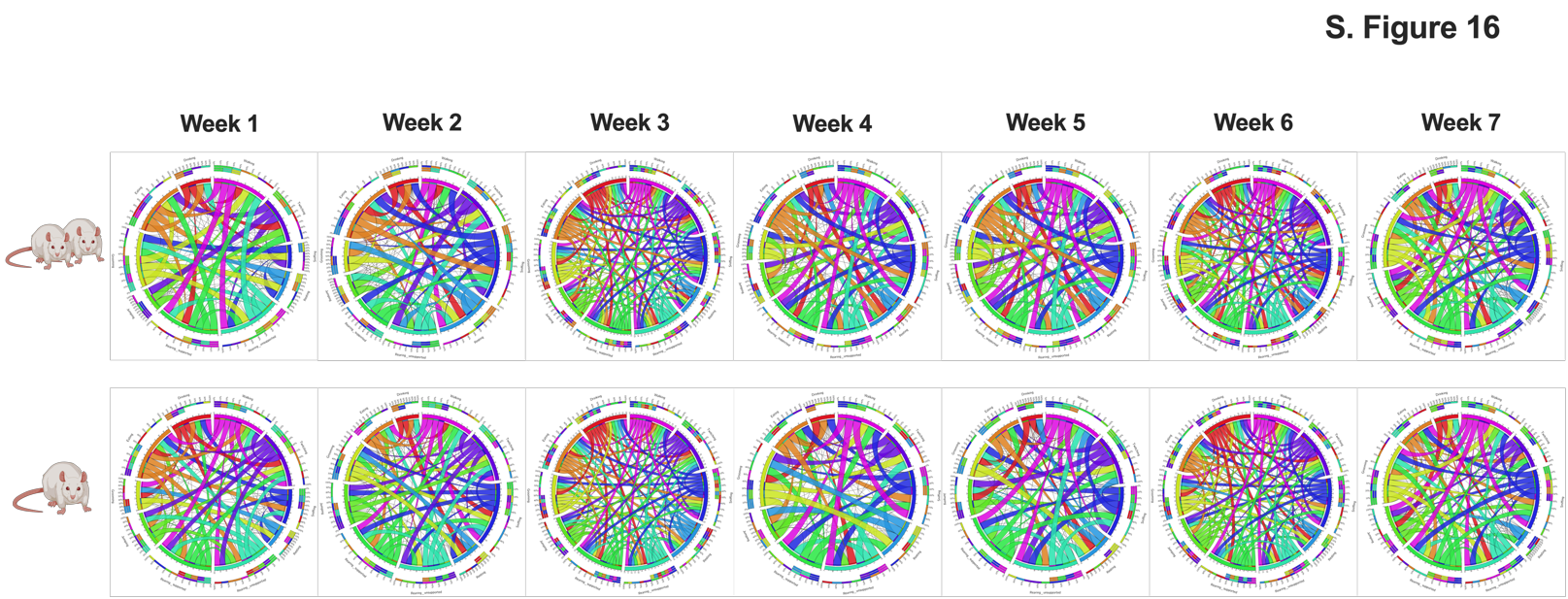
**S. Fig. 16.** *Organism-level behavioral associations throughout adolescence.* Connectograms display the change in behavioral associations for each behavior throughout the study. Paired animals are displayed on the top row, Isolated on the bottom. Thicker ribbons suggest strong associations.

**1.**

**10**

**9.**

**8.**

**7.**

**6.**

**5.**

**4.**

**3.**

**2.**

1. Walking
2. Twitching
3. Sniffing
4. Resting
5. Rearing Unsupported
6. Rearing Supported
7. Jumping
8. Grooming
9. Eating
10. Drinking

| **Behavior** | **Pairwise Comparisons** |  |
| --- | --- | --- |
| Adolescent Phenotypic Z Scores | t=3.218, df=22 *p*=0.0040 |  |
| Nonsignificant Behaviors | t=0.7506, df=22 *p*=0.4608 |  |
| Significant Behaviors | t=4.848, df=22 *p*<0.0001 |  |
|  | **2WAY ANOVA and Tukey HSD** |  |
| **Phenotypic Scores - Paired vs Isolated** |  | **Week** |
| Week 1 | *p*=0.9865 | *F (3.064, 63.33) = 48.97 p*<0.0001 |
| Week 2 | *p*=0.0784 |  |
| Week 3 | *p*=0.1764 | **Stress** |
| Week 4 | *p*=0.0041 | *F (1, 22) = 15.91 p* =0.0006 |
| Week 5 | *p*=0.0149 |  |
| Week 6 | *p*=0.0129 | **Week x Stress** |
| Week 7 | *p*=0.3726 | F (6, 124) = 5.388 p <0.0001 |

**S. Table 1. Phenotypic Z-Scores**

**S. Table 2. PC1 and PC2 Scores**

| **Week** | **Pairwise Comparisons** |
| --- | --- |
| **PC 1 Scores** |  |
| Week 1 | t=0.3988, df=22, 0.6939 |
| Week 2 | t=2.312, df=22, 0.0305 |
| Week 3 | t=1.233, df=22, 0.2307 |
| Week 4 | t=2.886, df=22, 0.0086 |
| Week 5 | t=2.145, df=22, 0.0433 |
| Week 6 | t=0.4405, df=22, 0.6639 |
| Week 7 | t=0.1946, df=22, 0.8475 |
| **PC 2 Scores** |  |
| Week 1 | t=4.497, df=22, 0.0002 |
| Week 2 | t=1.022, df=22, 0.3180 |
| Week 3 | t=2.394, df=22, 0.0256 |
| Week 4 | t=0.4375, df=22, 0.6660 |
| Week 5 | t=1.017, df=22, 0.3201 |
| Week 6 | t=3.379, df=22, 0.0027 |
| Week 7 | t=0.5471, df=22, 0.5898 |

**S. Table 3. Weekly Behaviors.**

| **Behaviors** | **Weekly** | **Stress** | **Interaction** |
| --- | --- | --- | --- |
| Eating | F (6, 146) = 2.865 *p*=0.0114 | F (1, 146) = 1.208 *p*=0.2736 | F (6, 146) = 0.4740 *p*=0.8268 |
| Drinking | F (6, 146) = 4.279 *p*=0.0005 | F (1, 146) = 0.6477 *p*=0.4223 | F (6, 146) = 0.1239 *p*=0.9933 |
| Grooming | F (6, 146) = 21.62 *p*<0.0001 | F (1, 146) = 0.0002278 *p*=0.9880 | F (6, 146) = 0.2157 *p*=0.9713 |
| Jumping | F (6, 146) = 19.99 *p*<0.0001 | F (1, 146) = 65.83 *p*<0.0001 | F (6, 146) = 3.169 *p*=0.0059 |
| Rearing Supported | F (6, 146) = 32.67 *p*<0.0001 | F (1, 146) = 0.2944 *p*=0.5882 | F (6, 146) = 1.962 *p*=0.0747 |
| Rearing Unsupported | F (6, 146) = 5.017 *p*=0.0001 | F (1, 146) = 6.797 *p*=0.0101 | F (6, 146) = 1.020 *p*=0.4148 |
| Resting | F (6, 146) = 0.8958 *p*=0.4998 | F (1, 146) = 1.694 *p*=0.1951 | F (6, 146) = 0.3960 *p*=0.8807 |
| Sniffing | F (6, 146) = 15.29 *p*<0.0001 | F (1, 146) = 18.90 *p*<0.0001 | F (6, 146) = 1.157 *p*=0.3328 |
| Twitching | F (6, 146) = 7.920 *p*<0.0001 | F (1, 146) = 68.16 *p*<0.0001 | F (6, 146) = 1.623 *p*=0.1448 |
| Walking | F (6, 146) = 19.50 *p*<0.0001 | F (1, 146) = 0.01932 *p*=0.8897 | F (6, 146) = 1.379 *p*=0.2267 |

**S. Table 4. Hourly Behaviors.**

| Behaviors | Interactions | Hours | Stress |
| --- | --- | --- | --- |
| Week 1 Drinking | F (12, 258) = 0.5402 P=0.8873 | F (12, 258) = 5.384 P<0.0001 | F (1, 258) = 1.431 P=0.2326 |
| Week 2 Drinking | F (12, 286) = 2.172 P=0.0131 | F (12, 286) = 7.175 P<0.0001 | F (1, 286) = 0.08270 P=0.7739 |
| Week 3 Drinking | F (12, 234) = 0.5940 P=0.8462 | F (12, 234) = 3.876 P<0.0001 | F (1, 234) = 0.4226 P=0.5163 |
| Week 4 Drinking | F (12, 286) = 0.8998 P=0.5476 | F (12, 286) = 6.136 P<0.0001 | F (1, 286) = 0.09116 P=0.7629 |
| Week 5 Drinking | F (12, 286) = 0.9431 P=0.5041 | F (12, 286) = 8.122 P<0.0001 | F (1, 286) = 0.3793 P=0.5385 |
| Week 6 Drinking | F (12, 234) = 0.5588 P=0.8736 | F (12, 234) = 3.253 P=0.0002 | F (1, 234) = 0.4332 P=0.5111 |
| Week 7 Drinking | F (12, 286) = 1.009 P=0.4402 | F (12, 286) = 5.508 P<0.0001 | F (1, 286) = 0.01138 P=0.9151 |
| Week 1 Eating | F (12, 258) = 0.9126 P=0.5349 | F (12, 258) = 5.312 P<0.0001 | F (1, 258) = 0.1521 P=0.6969 |
| Week 2 Eating | F (12, 286) = 1.463 P=0.1376 | F (12, 286) = 5.654 P<0.0001 | F (1, 286) = 0.3940 P=0.5307 |
| Week 3 Eating | F (12, 234) = 0.7173 P=0.7340 | F (12, 234) = 3.732 P<0.0001 | F (1, 234) = 0.8291 P=0.3635 |
| Week 4 Eating | F (12, 286) = 0.4968 P=0.9160 | F (12, 286) = 8.109 P<0.0001 | F (1, 286) = 0.1769 P=0.6744 |
| Week 5 Eating | F (12, 286) = 0.2643 P=0.9939 | F (12, 286) = 8.681 P<0.0001 | F (1, 286) = 2.723 P=0.1000 |
| Week 6 Eating | F (12, 234) = 0.1475 P=0.9997 | F (12, 234) = 5.323 P<0.0001 | F (1, 234) = 0.4774 P=0.4903 |
| Week 7 Eating | F (12, 286) = 1.454 P=0.1411 | F (12, 286) = 11.07 P<0.0001 | F (1, 286) = 1.688 P=0.1949 |
| Week 1 Grooming | F (12, 258) = 1.239 P=0.2564 | F (12, 258) = 7.648 P<0.0001 | F (1, 258) = 0.4605 P=0.4980 |
| Week 2 Grooming | F (12, 286) = 1.219 P=0.2693 | F (12, 286) = 4.557 P<0.0001 | F (1, 286) = 1.556 P=0.2133 |
| Week 3 Grooming | F (12, 234) = 1.002 P=0.4475 | F (12, 234) = 3.430 P=0.0001 | F (1, 234) = 0.8444 P=0.3591 |
| Week 4 Grooming | F (12, 286) = 0.5908 P=0.8492 | F (12, 286) = 4.950 P<0.0001 | F (1, 286) = 0.05130 P=0.8210 |
| Week 5 Grooming | F (12, 286) = 0.7158 P=0.7358 | F (12, 286) = 4.987 P<0.0001 | F (1, 286) = 0.5084 P=0.4764 |
| Week 6 Grooming | F (12, 234) = 0.4618 P=0.9352 | F (12, 234) = 3.078 P=0.0005 | F (1, 234) = 2.267 P=0.1335 |
| Week 7 Grooming | F (12, 286) = 1.045 P=0.4079 | F (12, 286) = 5.900 P<0.0001 | F (1, 286) = 0.4358 P=0.5097 |
| Week 1 Jumping | F (12, 258) = 1.284 P=0.2279 | F (12, 258) = 2.456 P=0.0048 | F (1, 258) = 17.35 P<0.0001 |
| Week 2 Jumping | F (12, 286) = 2.730 P=0.0016 | F (12, 286) = 3.863 P<0.0001 | F (1, 286) = 53.21 P<0.0001 |
| Week 3 Jumping | F (12, 234) = 0.8427 P=0.6063 | F (12, 234) = 1.251 P=0.2490 | F (1, 234) = 35.38 P<0.0001 |
| Week 4 Jumping | F (12, 286) = 1.235 P=0.2584 | F (12, 286) = 5.000 P<0.0001 | F (1, 286) = 48.03 P<0.0001 |
| Week 5 Jumping | F (12, 286) = 1.879 P=0.0366 | F (12, 286) = 9.750 P<0.0001 | F (1, 286) = 50.16 P<0.0001 |
| Week 6 Jumping | F (12, 234) = 1.653 P=0.0784 | F (12, 234) = 6.724 P<0.0001 | F (1, 234) = 37.69 P<0.0001 |
| Week 7 Jumping | F (12, 286) = 2.407 P=0.0055 | F (12, 286) = 11.86 P<0.0001 | F (1, 286) = 18.93 P<0.0001 |
| Week 1 Rearing | F (12, 258) = 0.3860 P=0.9678 | F (12, 258) = 13.40 P<0.0001 | F (1, 258) = 0.2080 P=0.6487 |
| Week 2 Rearing | F (12, 286) = 0.9270 P=0.5201 | F (12, 286) = 10.40 P<0.0001 | F (1, 286) = 2.423 P=0.1207 |
| Week 3 Rearing | F (12, 234) = 1.306 P=0.2155 | F (12, 234) = 6.049 P<0.0001 | F (1, 234) = 11.03 P=0.0010 |
| Week 4 Rearing | F (12, 286) = 0.6445 P=0.8033 | F (12, 286) = 14.74 P<0.0001 | F (1, 286) = 3.206 P=0.0744 |
| Week 5 Rearing | F (12, 286) = 0.6247 P=0.8208 | F (12, 286) = 8.258 P<0.0001 | F (1, 286) = 2.376 P=0.1243 |
| Week 6 Rearing | F (12, 234) = 0.5550 P=0.8764 | F (12, 234) = 9.158 P<0.0001 | F (1, 234) = 7.748 P=0.0058 |
| Week 7 Rearing | F (12, 286) = 1.016 P=0.4342 | F (12, 286) = 7.233 P<0.0001 | F (1, 286) = 2.096 P=0.1488 |
| Week 1 Rearing Unsp | F (12, 258) = 1.393 P=0.1692 | F (12, 258) = 4.468 P<0.0001 | F (1, 258) = 0.2946 P=0.5878 |
| Week 2 Rearing Unsp | F (12, 286) = 1.679 P=0.0707 | F (12, 286) = 5.630 P<0.0001 | F (1, 286) = 3.452 P=0.0642 |
| Week 3 Rearing Unsp | F (12, 234) = 1.078 P=0.3793 | F (12, 234) = 5.494 P<0.0001 | F (1, 234) = 14.23 P=0.0002 |
| Week 4 Rearing Unsp | F (12, 286) = 0.4985 P=0.9149 | F (12, 286) = 11.48 P<0.0001 | F (1, 286) = 2.523 P=0.1133 |
| Week 5 Rearing Unsp | F (12, 286) = 0.3993 P=0.9633 | F (12, 286) = 9.995 P<0.0001 | F (1, 286) = 6.526 P=0.0111 |
| Week 6 Rearing Unsp | F (12, 234) = 0.7520 P=0.6994 | F (12, 234) = 6.965 P<0.0001 | F (1, 234) = 0.01122 P=0.9157 |
| Week 7 Rearing Unsp | F (12, 286) = 1.049 P=0.4037 | F (12, 286) = 6.698 P<0.0001 | F (1, 286) = 0.03316 P=0.8556 |
| Week 1 Resting | F (12, 258) = 1.515 P=0.1186 | F (12, 258) = 2.558 P=0.0033 | F (1, 258) = 1.560 P=0.2128 |
| Week 2 Resting | F (12, 286) = 0.6619 P=0.7874 | F (12, 286) = 1.473 P=0.1337 | F (1, 286) = 5.572 P=0.0189 |
| Week 3 Resting | F (12, 234) = 0.4584 P=0.9369 | F (12, 234) = 1.058 P=0.3966 | F (1, 234) = 0.1583 P=0.6911 |
| Week 4 Resting | F (12, 286) = 0.4019 P=0.9623 | F (12, 286) = 3.018 P=0.0005 | F (1, 286) = 0.03398 P=0.8539 |
| Week 5 Resting | F (12, 286) = 0.5771 P=0.8602 | F (12, 286) = 2.865 P=0.0010 | F (1, 286) = 0.006831 P=0.9342 |
| Week 6 Resting | F (12, 234) = 1.072 P=0.3847 | F (12, 234) = 2.194 P=0.0127 | F (1, 234) = 5.160 P=0.0240 |
| Week 7 Resting | F (12, 286) = 1.235 P=0.2585 | F (12, 286) = 3.105 P=0.0004 | F (1, 286) = 0.07661 P=0.7821 |
| Week 1 Sniffing | F (12, 258) = 0.5875 P=0.8517 | F (12, 258) = 6.344 P<0.0001 | F (1, 258) = 0.04524 P=0.8317 |
| Week 2 Sniffing | F (12, 286) = 1.791 P=0.0492 | F (12, 286) = 6.370 P<0.0001 | F (1, 286) = 14.61 P=0.0002 |
| Week 3 Sniffing | F (12, 234) = 1.555 P=0.1058 | F (12, 234) = 4.052 P<0.0001 | F (1, 234) = 34.44 P<0.0001 |
| Week 4 Sniffing | F (12, 286) = 0.5400 P=0.8877 | F (12, 286) = 9.700 P<0.0001 | F (1, 286) = 16.44 P<0.0001 |
| Week 5 Sniffing | F (12, 286) = 0.9258 P=0.5214 | F (12, 286) = 7.295 P<0.0001 | F (1, 286) = 16.82 P<0.0001 |
| Week 6 Sniffing | F (12, 234) = 0.5672 P=0.8673 | F (12, 234) = 6.535 P<0.0001 | F (1, 234) = 1.640 P=0.2017 |
| Week 7 Sniffing | F (12, 286) = 1.158 P=0.3133 | F (12, 286) = 7.985 P<0.0001 | F (1, 286) = 1.928 P=0.1661 |
| Week 1 Twitching | F (12, 257) = 1.961 P=0.0282 | F (12, 257) = 10.02 P<0.0001 | F (1, 257) = 23.27 P<0.0001 |
| Week 2 Twitching | F (12, 286) = 2.071 P=0.0188 | F (12, 286) = 5.490 P<0.0001 | F (1, 286) = 25.74 P<0.0001 |
| Week 3 Twitching | F (12, 234) = 1.296 P=0.2216 | F (12, 234) = 3.491 P<0.0001 | F (1, 234) = 40.47 P<0.0001 |
| Week 4 Twitching | F (12, 286) = 1.042 P=0.4101 | F (12, 286) = 6.346 P<0.0001 | F (1, 286) = 19.56 P<0.0001 |
| Week 5 Twitching | F (12, 286) = 1.039 P=0.4132 | F (12, 286) = 7.242 P<0.0001 | F (1, 286) = 37.49 P<0.0001 |
| Week 6 Twitching | F (12, 234) = 1.316 P=0.2099 | F (12, 234) = 9.735 P<0.0001 | F (1, 234) = 10.84 P=0.0011 |
| Week 7 Twitching | F (12, 286) = 1.651 P=0.0773 | F (12, 286) = 8.310 P<0.0001 | F (1, 286) = 3.481 P=0.0631 |
| Week 1 Walking | F (12, 258) = 0.4536 P=0.9396 | F (12, 258) = 7.709 P<0.0001 | F (1, 258) = 3.666 P=0.0566 |
| Week 2 Walking | F (12, 286) = 1.677 P=0.0712 | F (12, 286) = 10.07 P<0.0001 | F (1, 286) = 2.356 P=0.1259 |
| Week 3 Walking | F (12, 234) = 1.148 P=0.3224 | F (12, 234) = 4.008 P<0.0001 | F (1, 234) = 9.146 P=0.0028 |
| Week 4 Walking | F (12, 286) = 0.6495 P=0.7988 | F (12, 286) = 12.47 P<0.0001 | F (1, 286) = 1.620 P=0.2041 |
| Week 5 Walking | F (12, 286) = 0.5671 P=0.8679 | F (12, 286) = 6.471 P<0.0001 | F (1, 286) = 1.577 P=0.2103 |
| Week 6 Walking | F (12, 234) = 0.5562 P=0.8755 | F (12, 234) = 8.588 P<0.0001 | F (1, 234) = 10.05 P=0.0017 |
| Week 7 Walking | F (12, 286) = 0.9182 P=0.5290 | F (12, 286) = 6.178 P<0.0001 | F (1, 286) = 1.479 P=0.2249 |

**S. Table 5. Residual Avoidance. Two-Way ANOVA.**

| **Behavior** | **Weeks** | **Stress** | **Interaction** |
| --- | --- | --- | --- |
| Shelter Zone | F (6, 141) = 1.511 *p*=0.1786 | F (1, 141) = 2.283 *p*=0.1330 | F (6, 141) = 1.511 *p*=0.1786 |
| Food Zone | F (6, 141) = 0.7140 *p*=0.6389 | F (1, 146) = 1.461 *p*=0.0470 | F (6, 141) = 0.7140 *p*=0.6389 |
| Food Zone during spotlight | F (6, 141) = 0.9163  *p*=0.4852 | F (1, 141) = 7.545  *p*=0.0068 | F (6, 141) = 0.6354  *p*=0.7017 |

**S. Table 6. Acoustic Startle Reactivity**

| **Behavior** | **Pairwise Comparison** |
| --- | --- |
| mV_Max | t=1.974, df=20 *p*=0.0624 |
| T_Max | t=0.9167, df=21 *p*=0.3697 |
| mV_AVG | t=2.040, df=20 *p*=0.0548 |
| **mV_Max** |  |
| Block 1 | t=1.777, df=20 *p*=0.0908 |
| Block 2 | t=0.9544, df=21 *p*=0.3508 |
| Block 3 | t=1.484, df=20, *p*-0.1535 |
| **mV_AVG** |  |
| Block 1 | t=1.527, df=20 *p*= 0.1424 |
| Block 2 | t=0.9202, df=21 *p*=0.3679 |
| Block 3 | t=1.370, df=21 *p*=0.1853 |

**S. Table 7. Elevated Plus Maze**

| **Behavior** | **Pairwise Comparison** |
| --- | --- |
| Anxiety Index | t=1.070, df=22 *p*=0.2961 |
| **CA** |  |
| Time | t=1.174, df=21 *p*=0.2535 |
| Frequency | t=1.213, df=22 *p*=0.2380 |
| **OA** |  |
| Time | t=1.174, df=21 *p*=0.2535 |
| Frequency | t=1.213, df=22 *p*=0.2380 |
| **Body Elongation Stretch** |  |
| Frequency | t=2.224, df=22 *p*=0.0367 |
| Distanced moved (cm) | t=0.7794, df=22 *p*=0.4441 |

**S. Table 8. Social Y Maze**

| **Behavior** | **Pairwise Comparison** |
| --- | --- |
| **Object** |  |
| Frequency | Sum of Ranks (127, 104), *p*=0.6873 |
| CD | t=0.08243, df=22 *p*=0.9350 |
| **Conspecific** |  |
| Frequency | t=1.392, df=22 *p*=0.1779 |
| CD | t=2.143, df=22 *p*=0.0434 |
| **Body Elongation Stretch** |  |
| Frequency | t=3.467, df=22 *p*=0.0022 |
| CD | Sum of Ranks (161, 49), *p*<0.0001 |
| **Distance Traveled** | t=1.934, df=21 *p*=0.0667 |

**S. Table 9. Emotionality**

| **Behavior** | **Student t** |
| --- | --- |
| **SYM Z Norm** |  |
| Z Norm Conspecific Duration | t=2.143, df=22 *p*=0.0434 |
| Z Norm SAP Frequency | t=3.467, df=22 *p*=0.0022 |
| Z Norm Object Duration | t=0.08243, df=22 *p*=0.9350 |
| **EPM Z Norm** |  |
| Z Norm CA Frequency | t=0.07935, df=22 *p*=0.9375 |
| Z Norm Latency to CA | t=0.04545, df=22 *p*=0.9642 |
| Z Norm SAP Duration | t=2.415, df=22 *p*=0.0245 |
| **ASR** |  |
| Z Norm mV_AVG | t=2.040, df=20 *p*=0.0548 |
| Z Norm T_Max Block 1 | t=2.314, df=21 *p*=0.0309 |
| Z Norm mV_Max | t=1.974, df=20 *p*=0.0624 |
| **Emotionality** | t=4.017, df=22 *p*=0.0006 |
| SYM Z Avg | t=2.139, df=20 *p*=0.0449 |
| EPM Z Avg | t=1.008, df=22 *p*=0.3243 |
| ASR Z Avg | t=3.693, df=22 *p*=0.0013 |

**S. Table 10. Divergent Sex Effects**

|  | **Two-Way ANOVA and Tukey HSD Post Hoc Comparisons** |
| --- | --- |
| Interaction | F (1, 61) = 9.032 *p*=0.0038 |
| Sex | F (1, 61) = 33.85 *p*<0.0001 |
| Stress | F (1, 61) = 1.503 *p*=0.2250 |
| Males Paired v Males Isolated | *p*=0.5985 |
| Females Paired v Females Isolated | *p*=0.0191 |

**S. Table 11. Binge-Like Eating and Correlations**

| BED Hyperphagic | t=3.588, df=9 *p*=0.005 | Pairwise comparisons |
| --- | --- | --- |
| Twitching and WD consumption | r= 0.5552, p=0.0049, n=24 | Pearson correlation |
| Jumping and WD consumption | r= 0.5150, p=0.0102, n=24 | Pearson correlation |
| Hyperphagic and hypophagic twitching comparisons | Interaction: F (1,20)=3.063, *p*=0.0954 Binge-like Effect: F (1,20) =5.032, *p*=0.0364 Stress Effect: F (1,20) =35.28, *p*<0001 Hyperphagic:Paired vs. Hyperphagic:Isolated, *p*=0.0002; Hyperphagic:Paired vs. Hypophagic:Paired, *p*=0.9847; Hyperphagic:Paired vs. Hypophagic:Isolated, *p*=0573; Hyperphagic:Isolated vs. Hypophagic:Paired, *p*=0.0001; Hyperphagic:Isolated vs. Hypophagic:Isolated, *p*=0.0495; Hypophagic:Paired vs. Hypophagic:Isolated, *p*=0.0269 | Two-Way ANOVA with Tukey HSD post hoc |
|  | Paired | Isolated |
| Jumping and BED Paired Isolated Separate | r= 0.047, p=0.884 | r= 0.570, p=0.053 |
| Twitching and BED Paired Isolated Separate | r= 0.265, p=0.405 | r= 0.657, p=0.020 |

**S. Table 12. Endocrine**

|  | Pairwise Comparisons |
| --- | --- |
| CORT | t=4.006, df=20 *p*=0.0007 |
| Estrogen | t=0.5395, df=22 *p*=0.5949 |
| Progesterone | t=1.280, df=18 *p*=0.2170 |
| LH | t=0.4650, df=22 *p*=0.6465 |
|  | Spearman Correlation |
| Adolescent Shifted Behavioral z-score and CORT | r=-0.428, *p*=0.042 |
| Emotionality and CORT | r=-0.418, *p*=0.0421 |
|  | Pearson Correlation |
| Jumping and Cort | r=-0.404, *p*=0.050 |
| Twitching and CORT | r=-0.463, *p*=0.023 |
